# Plasma membrane folate transport in fungi and plants is mediated by members of the oligopeptide transporter (OPT) family

**DOI:** 10.1101/2025.06.23.661034

**Authors:** Nikita Vashist, Md Shabbir Ahmad, Sahithi Vedula, Chinmayee Choudhury, Sunil Laxman, Anand K Bachhawat

## Abstract

Folates are essential for all organisms. They are acquired either through *de novo* biosynthesis or from the diet. Yeast, fungi and plants make their own folates and it has not been clear if plasma membrane folate transporters exist in these organisms. Using a synthetic lethal screen in *Saccharomyces cerevisiae* we observed that deletions in a gene encoding the previously identified glutathione transporter, OPT1, was synthetically sick with a disruption in folate biosynthesis. Uptake experiments confirmed that Opt1p/Hgt1p can transport folinic acid and the naturally abundant methyl tetrahydrofolate. As *S. cerevisiae* Opt1p was able to transport both folate and glutathione, we used alanine-scanning mutants of the residues in the transmembrane domains of the channel to identify the residues required specifically for the uptake of folates and distinct from those required for glutathione. We further examined the oligopeptide transporter family of other organisms for the presence of folate transporters. In *C. albicans*, CaOPT1, the orthologue of *S. cerevisiae* OPT1 efficiently transported folate but not glutathione, while the previously characterized glutathione transporter, CaOPT7 could not transport folate. *Aspergillus fumigatus* has eight homologues of the oligopeptide transporter family, of which OptB and OptH could transport folates. In the plant *Arabidopsis thaliana,* the Opt1 homologs AtOpt2, AtOpt4, and AtOpt6 could transport folates. This discovery of folate transporters across fungi and plants fills a critical gap in our understanding of folate metabolism, and can benefit the exploitation of these pathways in pathogenic fungi, and in plants.

## Introduction

Folates are essential metabolites in all living organisms. They facilitate the transfer of one-carbon units by acting as their donors in several metabolic pathways (Appling, 1991). They contribute to the production of nucleotides, which are essential for cell proliferation, growth, and repair. Folates also help regenerate methionine from homocysteine, supporting methylation reactions crucial for cell function. Additionally, folates facilitate the interconversion of serine and glycine, and are important in histidine catabolism (Lucock et al., 2002), and are vital for the formation of N-formylmethionyl-tRNA in all organisms, which aids in mitochondrial protein translation and function (Appling, 1991)

Mammals lack the folate biosynthetic pathway, and obtain folates through their diets. Two transmembrane carriers mediate cellular folate uptake in humans - SLC19A1(a reduced folate carrier), and SLC46A1 (a proton coupled folate transporter) (Zhao et al., 2011). Unlike mammals, other eukaryotes such as yeast, fungi and plants can synthesize folates *de novo*. Folates are tripartite molecules consisting of pterin, p-amino benzoic acid (PABA) and glutamate. The enzymes involved in the biosynthesis of these moieties leading to folate formation has been characterized in different organisms (Gorelova et al., 2019; Green and Matthews, 2007; Rébeillé et al., 2006). In *Saccharomyces cerevisiae*, pterins are synthesized from GTP through the action of the enzymes FOL2, FOL1 and FOL3. The pterin then links up with 4-aminobenzoate and L-glutamate moieties to form folates. The folates are eventually reduced to tetrahydrofolate (THF), before they enter the folate cycle (Lagosky et al., 1987). In yeasts and plants some steps of the pathway are also completed in the mitochondria and chloroplast, and some of the intermediates are transported across the subcellular membrane through specific transporters. However, transport of folates across the plasma membrane has not been known in these organisms.

In *S. cerevisiae*, disruption of any of the folate biosynthetic genes FOL2, FOL1, or FOL3 leads to folate auxotrophy. These deletions can still survive when external folates like folinic (5-formyl THF) acid or folic acid are added to the medium. The dependence of folate auxotrophs on external folates suggests the existence of a plasma membrane folate transporter in yeast (Cherest et al., 2000; Güldener et al., 2004; Little and Haynes, 1979; Nardese et al., 1996). Similarly, exogenous folates can stimulate the growth of wild type *Candida glabrata* (Porzoor and Macreadie, 2015), again suggesting the existence of folate transport mechanisms. There have therefore been several efforts to investigate and identify plasma membrane folate transporters in yeast. The efforts included bioinformatic approaches looking for homologs of the human folate transporters (Macreadie, 2016) as well as experimental approaches using antifolate inhibitors such as dihydropteroate (Bayly and Macreadie, 2002), sulfa drugs (Bayly et al., 2001) and methotrexate (Wong et al., 2017). However, none of these efforts have successfully identified the folate transporter. Thus, the plasma membrane folate transporter of yeasts and fungi has remained unknown.

In this study, we adopted a genetic approach to identify a folate transporter in the yeast *S. cerevisiae*. In this strategy we have exploited an expected synthetic lethality between the knockout of a candidate folate transporter and a knockout in the folate biosynthetic pathway. Through this, we successfully identify a plasma membrane folate transporter in yeast. We found, surprisingly, that the previously characterized glutathione transporter, Opt1p/Hgt1p, also effectively transported folates. Opt1p/Hgt1p is a member of the oligopeptide transporter family, whose members are found in yeast, plants, and fungi but not in mammals or metazoans. Most members of this oligopeptide transporter family have still not been assigned any function. We show that several of its homologs in other organisms, including pathogenic yeast, fungi, and plants functioned as folate transporters. While some transporters were exclusive to either glutathione or folate, some could transport both. As Opt1p/Hgt1p was one of the transporters that was able to transport both folate and glutathione, we further mapped the residues critical for folate transport using an alanine-scanned mutant library of the TMDs lining the channel pore. This study addresses a critical gap in folate metabolism, and has important implications in efforts to exploit these pathways in both pathogenic fungi and plants.

## Results

### Genetic screen to identify folate transporters using *Saccharomyces cerevisiae* identifies Opt1

The yeast *Saccharomyces cerevisiae* carries out *de novo* biosynthesis of folates through the sequential action of the enzymes encoded by FOL2, FOL1 and FOL3 genes. Deletion of any of these genes is lethal unless supplemented with folates. We used these observations to develop a synthetic lethal screen to identify a folate transporter in yeast. We first examined the database of *Saccharomyces cerevisiae* for possible synthetic lethal hits with either *fol2*Δ*, fol1*Δ *or fol3*Δ since many genome-wide synthetic lethal screens have been carried out (SGD, https://www.yeastgenome.org). However, we did not identify any hits from data mining.

Using a *fol2*Δ*::HIS3* disruption cassette, we evaluated a library of knockouts of uncharacterized/putative transporter strains, obtained from EUROSCARF. The transporter deletions were transformed with a linearized *fol2*Δ*::HIS3* cassette that had 496bp and 657bp of the FOL2 gene flanking the HIS3 cassette after excision from a plasmid. The transformants were selected on minimal media containing folinic acid and were screened for growth in media lacking folinic acid. In the case of transporters that were not folate transporters, we expected to see many colonies that would show folinic acid auxotrophy. However, if the transporter deletion was a folate transporter, we would expect minimal colonies, with none exhibiting folinic acid auxotrophy. In the latter case, the few His^+^ colonies that would arise, would be due to the cassette being integrated at some other locus rather than the FOL2 locus. This was because the *fol2*Δ along with folate transporter deletion would be either sick or lethal due to folate depletion resulting from the loss of both endogenous folate biosynthesis and the ability to import folates from the environment. The detailed experimental strategy to identify a folate transporter in *S. cerevisiae* is summarized in Fig.1A.

**Fig.1:**
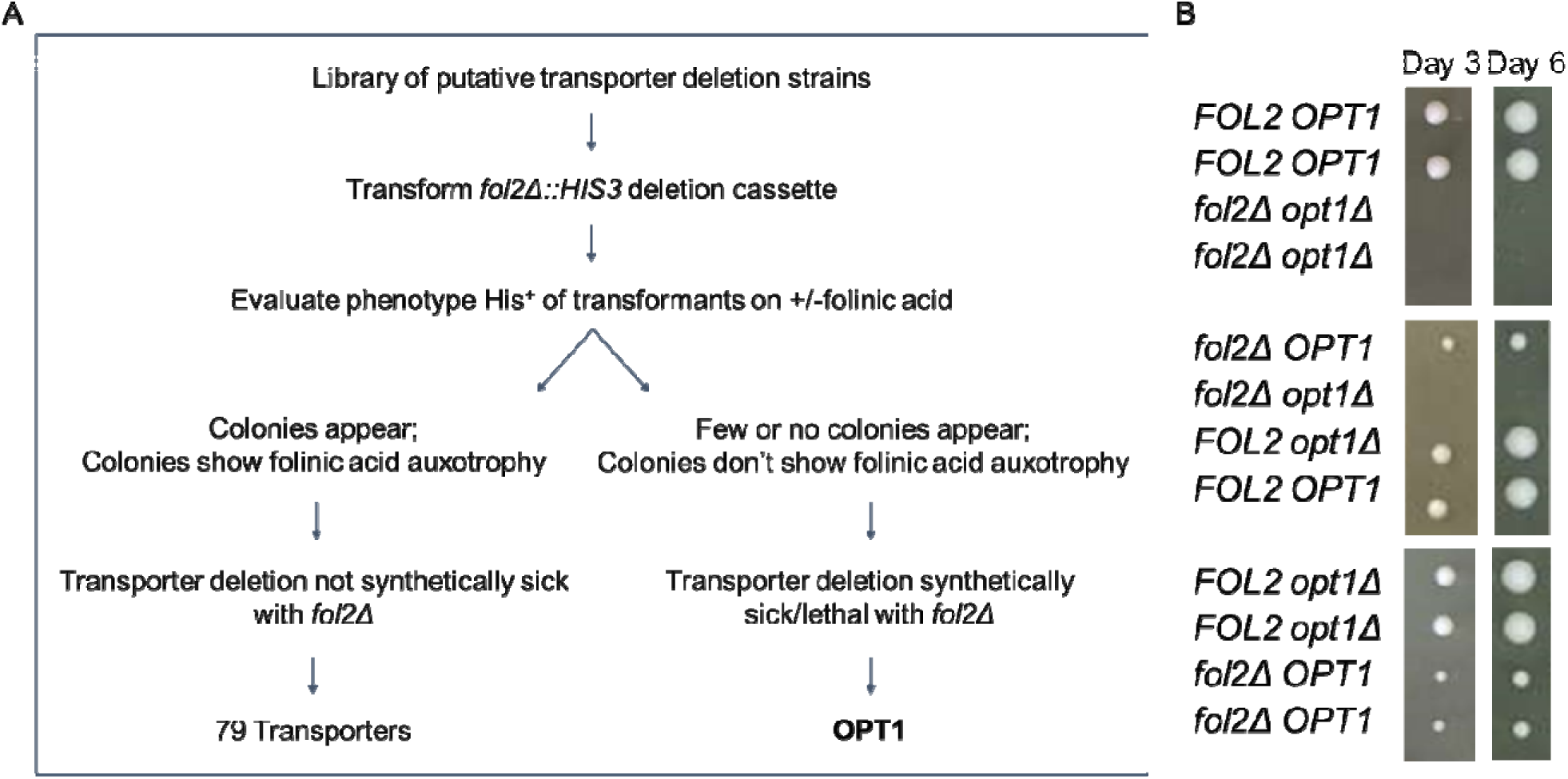
*fol2*Δ *opt1*Δ exhibits a severe growth defect. A. Flowchart depicting the experimental strategy to identify a folate transporter in *S. cerevisiae.* A synthetic lethal screen was performed by transforming the *fol2*Δ::HIS3 disruption cassette i a knockout library of putative transporters from EUROSCARF. Transformants were selected on minimal media with folinic acid and screened for growth on media without folinic acid. Non-folate transporters strains were expected to show folinic acid auxotrophy, whereas folate transporter deletion along with *fol2*Δ would result in fewer colonies, with no folinic acid auxotrophy. B. Tetrad analysis of the heterozygous diploid strain W303 *fol2*Δ */ FOL2 opt1*Δ */ OPT1* strain was carried out. The tetrads were evaluated on YPAD medium supplemented with 100μM folinic acid. Plates were incubated at 30°C for upto 6 days. Photographs were taken on third and sixth day of incubation at 30 °C.

Strains carrying deletions in different transporters were procured from EUROSCARF and were screened using this approach. Among the 80 strains screened, 79 yielded disruptants exhibiting folinic acid auxotrophy. Only one strain with a deletion in the OPT1 transporter failed to yield *fol2* disruptants. OPT1, also known as HGT1, belongs to the Oligopeptide Transporter (OPT) family and encodes a plasma membrane glutathione transporter (Bourbouloux et al., 2000). To examine the potential role of OPT1 in folate transport more rigorously, we repeated the deletion with the same cassette looking at the transformant colonies in the *opt1*Δ versus the WT background in greater numbers (Supplementary Table S1). It was observed that, in the wild-type strain, the majority of His transformants were auxotrophic for folinic acid, whereas in the *opt1*Δ strain, there were significantly fewer transformants, and none displayed folinic acid auxotrophy (Supplementary Fig. S1). This confirmed that the double deletion was not being created as it was either very sick or lethal.

### Genetic analysis reveals that the *fol2***Δ** shows a synthetically sick phenotype with *opt1***Δ**

The inability to create a *fol2*Δ in an *opt1*Δ background suggested that the double deletion was either lethal or very sick. To examine this rigorously, we decided to construct and evaluate the double deletion *fol2*Δ*opt1*Δ using the classic genetic approach of tetrad dissection. Since *S. cerevisiae* BY4741 strains are known to be poor sporulators, we evaluated this using the *S. cerevisiae* W303 background. *fol2*Δ and *opt1*Δ were each created in the opposite mating types of W303, crossed and sporulated and the tetrads were evaluated. A total of 32 tetrads were obtained (Supplementary Table S2) and a representative image of the different non-parental, parental and tetratype tetrads is shown in Fig.1B. Among the four types of spores, all *fol2*Δ spores showed a slightly slower growth. This is because even exogenously added folate does not restore growth of the folate auxotrophs fully as compared to WT (Güldener et al., 2004; Nardese et al., 1996). After 3 days all colonies grew except the *fol2*Δ *opt1*Δ. However, when we prolonged the incubation upto 6 days, tiny colonies became visible but no further growth was observed. The severe growth defect of *fol2*Δ *opt1*Δ implies that Opt1p which is otherwise known to be the sole glutathione transporter *in S. cerevisiae* may also be the primary folate transporter in yeast. As the presence of glutathione (GSH) in yeast extract, component of YPAD medium, may also interfere with the growth (see following sections), we also dropped the spores from the tetrads directly on minimal medium supplemented with folinic acid. However, we observed very poor growth of the tetrads on this minimal medium, and no four-spore tetrads grew. We therefore continued to use the folinic acid supplemented YPAD medium for the tetrad dissection experiments.

### Growth of folate auxotrophs on folinic acid is inhibited by glutathione

Opt1p is the sole plasma membrane transporter of GSH in *S. cerevisiae* and has been very well characterized (Bourbouloux et al., 2000; Kaur and Bachhawat, 2009; Zulkifli et al., 2016). However, it also appeared as the primary transporter of folinic acid from the genetic analysis. We were interested to know, therefore, whether the two substrates interfered with each other. To investigate this, we supplemented the plates containing folinic acid with 200μM of GSH and examined the growth of *fol2*Δ strain. Despite the presence of folinic acid at 100μM, the *fol2*Δ strain failed to grow when the plates also contained 200μM GSH. This inhibitory effect was specific to GSH, and was not observed with other organic sulfur sources like methionine or cysteine (Fig.2).

**Fig.2:**
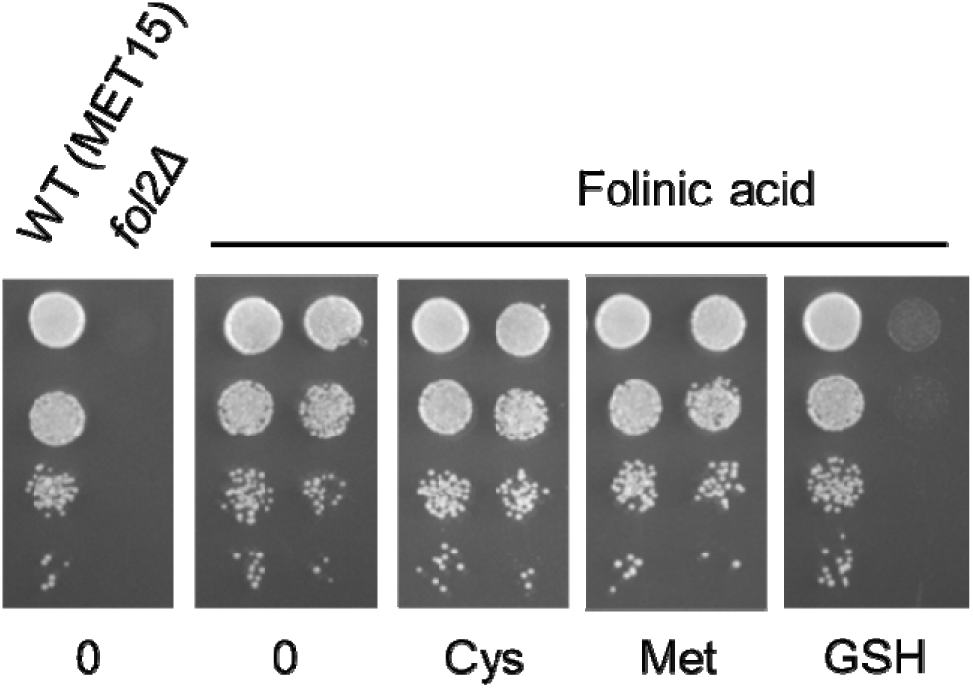
Glutathione inhibits utilization of folinic acid in the *fol2*Δ strain. The competition between GSH and folinic acid for uptake in the *fol2*Δ strain (BY4742 background) was evaluated using a growth assay. Plates contained SD medium supplemented with 200µM different sulfur source and 100µM folinic acid. The yeast nitrogen base used for the assay contains 2 μg/l of folic acid, which is insufficient for the growth of the *fol2*Δ strain.

To determine the concentration at which GSH begins to inhibit folate uptake, a range of GSH concentrations were tested. It was observed that 50µM GSH caused a reduction in growth, with nearly complete inhibition of growth occurring at 200µM (Supplementary Fig. S2)

We also investigated whether the reverse is true i.e. if folinic acid interferes with GSH utilization. To test this, we used a *met15*Δ strain, which is an organic sulfur auxotroph and thus depends on an organic sulfur source, in this case GSH, for growth. The strain was grown with 25μM GSH as the organic sulfur source along with increasing concentration of folinic acid. We observed that at 400µM of folinic acid the growth was completely inhibited (Supplementary Fig. S3). This suggests that GSH and folate each inhibit the utilization of the other substrate by Opt1p. This further confirms that they share the common transporter, Opt1p.

### Opt1p/Hgt1p can transport different folates

The ability of Opt1p to transport folates needed to be evaluated by uptake experiments. We therefore carried out uptake experiments, using specific detection by targeted liquid chromatography/mass spectrometry (LC-MS/MS) where the different folate forms could be distinguished (Supplmementary Table S3). We compared uptake in a *opt1*Δ strain transformed with either empty vector (control) or OPT1 expressed downstream of the constitutive TEF promoter. Upon addition of folinic acid, a rapid increase in intracellular folinic acid was observed in the cells bearing TEF-OPT1 plasmid, reaching saturation within 5 minutes (Fig.3A).

**Fig.3:**
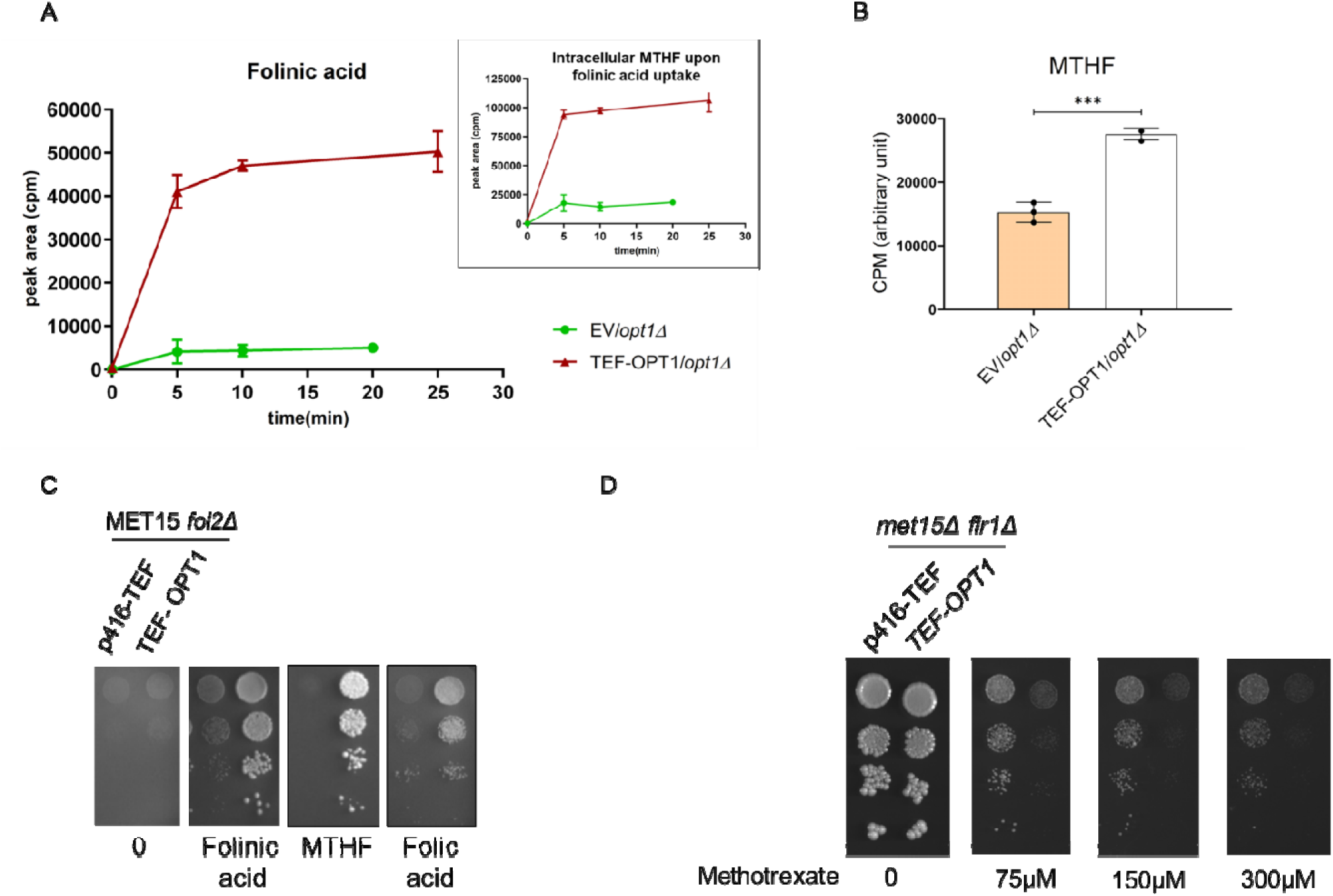
Opt1p transports different folate forms. A. Folinic acid uptake in yeast and its conversion to MTHF. Folinic acid uptake was estimated in an *opt1*Δ strain transformed with either vector control or TEF-OPT1 after 100µM folinic acid supplementation. Intracellular folate levels were measured at different time points. Folinic acid and MTHF were detected in the cells using targeted liquid chromatography/mass spectrometry (LC-MS/MS). Inset: MTHF levels in cells incubated with folinic acid. B. Uptake of MTHF by TEF-OPT1 MTHF uptake was estimated in an *opt1*Δ strain transformed with either vector control or TEF-OPT1 after 100µM MTHF supplementation. Intracellular MTHF levels was assessed after 5 minutes. Measurements were made using targeted liquid chromatography/mass spectrometry (LC-MS/MS). C. Ability of *S. cerevisiae* to grow on different forms of folate i.e. folinic acid, folic acid, and MTHF. Growth of *fol2*Δ strain transformed with vector control or TEF-OPT1 assessed on vitamin-free SD medium supplemented with approx. 50μM each of folinic acid, MTHF and folic acid. D. Overexpression of OPT1 increases sensitivity to methotrexate in *S. cerevisiae.* Growth of *flr1*Δ strain transformed with vector control or TEF-OPT1 assessed on vitamin free SD medium supplemented with methotrexate. Plates were incubated at 30°C for up to six days. Overexpression of OPT1 led to increased sensitivity to methotrexate across all tested concentrations, from 75 μM to 300 μM.

It is possible that the folinic acid that was taken up was being converted to other folates forms intracellularly (Supplementary Fig. S4). Therefore, in addition to examining folinic acid uptake, we also examined the predominant folate form, methyl tetrahydrofolate (MTHF). We observed that folinic acid conversion to MTHF was rapid (Fig.3A, inset). Owing to this rapid conversion of folinic acid to MTHF, determination of the absolute Km is difficult.

To assess whether OPT1 could also transport MTHF, the most abundant natural folate, we incubated MTHF with the *opt1*Δ cells transformed with vector or TEF-OPT1 and examined its uptake. Its levels were measured using LC-MS/MS at five minutes. We could observe that similar to folinic acid, MTHF was also effectively transported by Opt1p. (Fig.3B)

These experiments were also conducted in parallel on agar plates. In the plate experiments, instead of using an *opt1*Δ strain, we used a *fol2*Δ strain in a MET15 background (BY4742). These strains were either transformed with vector or TEF-OPT1 and assessed for growth on different folates. SD medium containing vitamin free yeast nitrogen base with methionine lacking folinic acid was used. Under these conditions, we could clearly see that folinic acid and MTHF were able to support the growth of *fol2*Δ strain expressed with OPT1 (Fig.3C) and supported the uptake experiment. Although we could not do the uptake experiment with folic acid owing to the low solubility, in plate growth experiments we clearly observe that it could also support the growth in an OPT1 dependent manner (Fig. 3C). Thus, all three folate forms tested i.e. folinic acid, MTHF and folic acid are transported by Opt1p.

### OPT1 overexpression leads to enhanced methotrexate sensitivity in *S. cerevisiae*

Methotrexate is an antifolate agent that competitively inhibits dihydrofolate reductase (DHFR), a key enzyme in the tetrahydrofolate biosynthetic pathway. Given the structural similarity between folate and methotrexate, it was hypothesized that Opt1p might facilitate the transport of both compounds.

Since Flr1 has been shown to mediate methotrexate efflux in *S. cerevisiae* (Brôco et al., 1999), a *flr1*Δ strain was used for evaluation. In this background, either a vector control or TEF-OPT1 was expressed, and growth was assessed on plates containing methotrexate. It was observed that overexpression of OPT1 led to increased sensitivity to methotrexate across all tested concentrations, from 75 μM to 300 μM (Fig. 3D).

### Identification of residues required for folate uptake in the Opt1p/Hgt1p transporter

The observation that Opt1p/Hgt1p transports both glutathione and the folates was probed further. This is because while folate is a tripartite molecule consisting of glutamate, PABA and pterin moieties, glutathione is a tripeptide consisting of glutamate, cysteine and glycine and is structurally quite distinct. We were therefore interested in comparing the recognition and transport of these two substrates, and to identify signatures that might uniquely identify folate transporters among the large family of oligopeptide transporters.

The ability of this transporter to transport glutathione has been previously extensively characterized through an alanine scanning mutagenesis of the entire 13 TMDs to identify the resides involved in binding and translocation of the substrate GSH. Since a structure was lacking, an *ab initio* model had previously been constructed to identify the TMDs forming the channel lining the pore (Zulkifli et al., 2016; Zulkifli and Bachhawat, 2017). The alanine scanning mutagenesis identified several residues in the TMDs 3,4,5,6,7,9,13 lining the channel. This *ab initio* model was further refined in the loop regions using the alphafold2 structure, and an energy grid was generated keeping the centroid of the TMD residues as grid centre. Then we docked both folinic acid and glutathione to this grid. The docking calculations revealed that folinic acid and glutathione bind to yeast OPT1 with the best docking scores of −6.72 and −4.26 respectively, indicating that they both do bind the transporter, and that it had better binding affinities of folinic acid over glutathione.

Using this alanine scanned mutant library of these TMDs, we sought to examine whether these libraries would enable us to identify which of these residues were important for folate recognition, as opposed to those required for glutathione. Some residues, those which were important for the protein expression, localization and translocation process, independent of substrate recognition, would of course be defective in both folate and glutathione transport. We individually transformed each of the alanine mutants (TMD by TMD) into the *met15*Δ *fol2*Δ strain, and the transformants were then spotted on SD medium using cysteine as a sulfur source along with limiting folinic acid. We used cysteine as a sulfur source as we found that it was most suitable for large scale screens (Supplementary Fig. 5) Methionine with folinic acid supplemented plates were used as a control. We carried out the growth experiments of the different TMD mutants and evaluated the results (Fig. 4). Only the TMDs that had residues that played a role were included in this figure and thus TMD13 is not included but depicted separately (Supplementary Fig. S6). These results are also summarized in a tabulated format (Table 1).

**Table 1.**
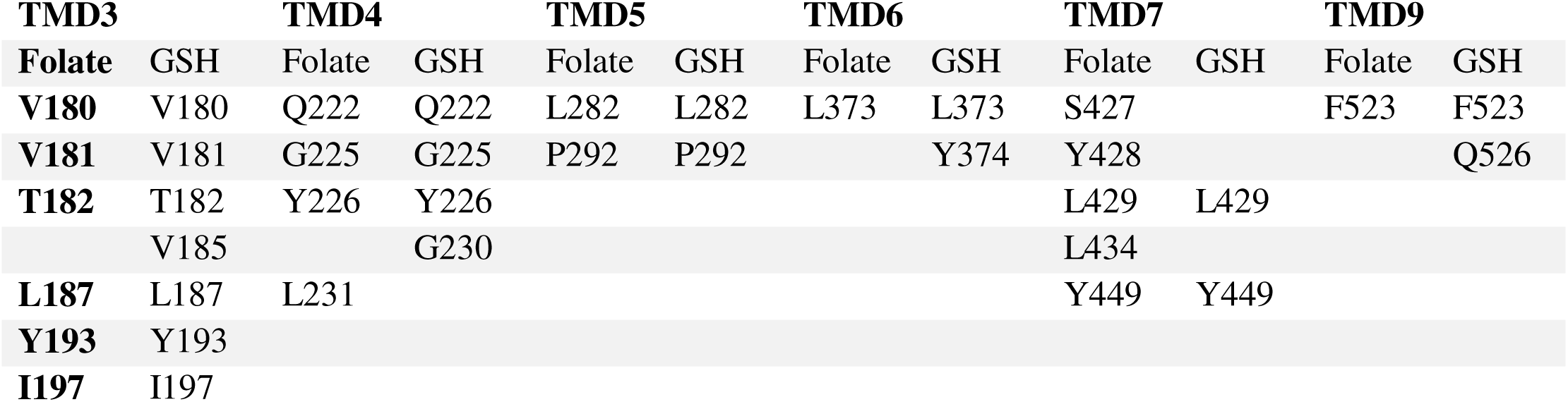
List of residues in TMD3, TMD4, TMD5, TMD7 and TMD9 essential for either folate transport or GSH transport based on the growth assay of the alanine mutants of these residue.

**Fig.4:**
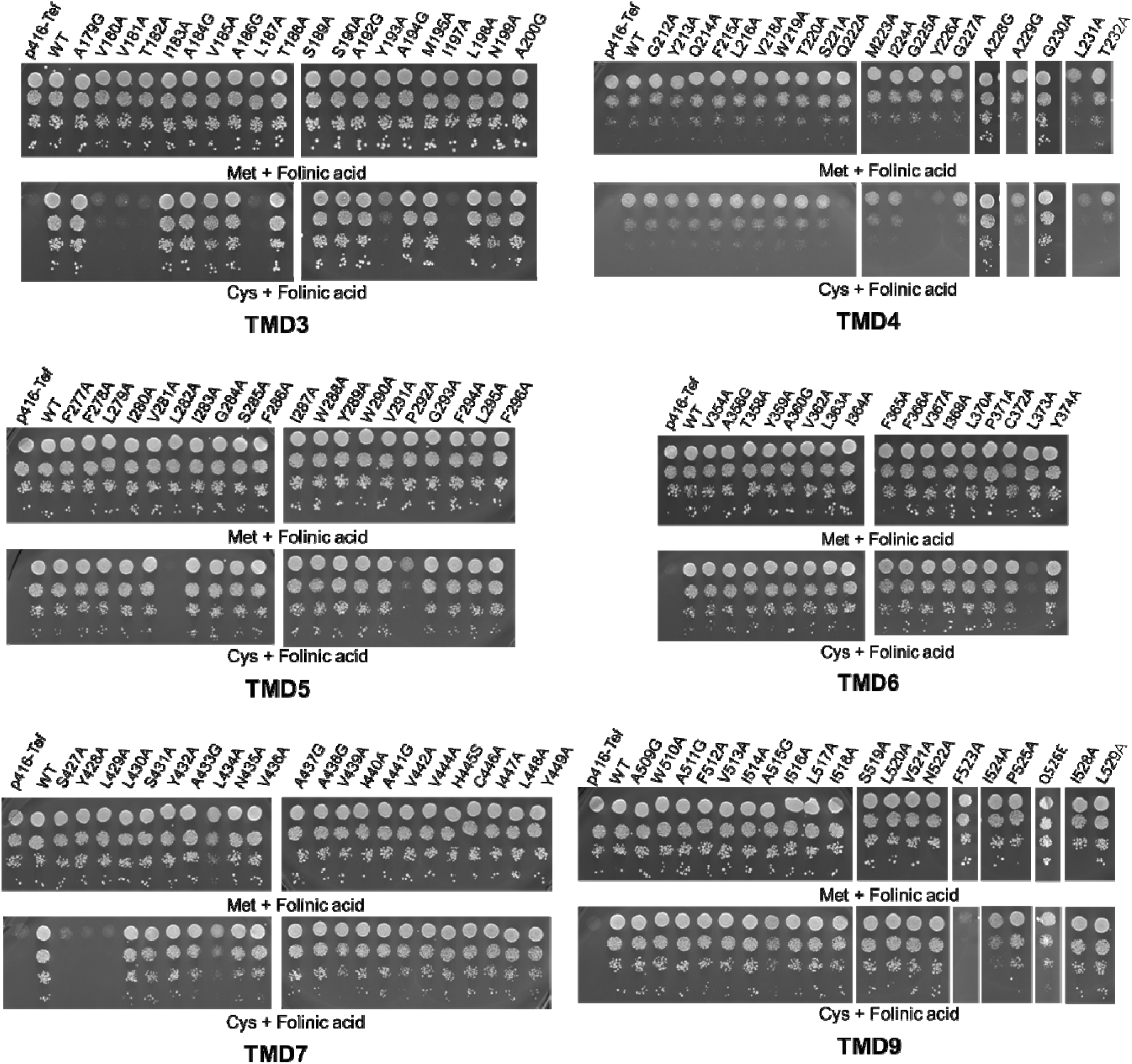
Functional characterization of alanine mutants of TMD3, 4, 5, 6, 7, 9 of Opt1p. OPT1/HGT1 and the different alanine mutants under the TEF promoter and corresponding vector (p416TEF) were transformed into strain *fol2*Δ *met15*Δ strain and evaluated for the ability to transport folinic acid using the growth assay by dilution spotting on minimal medium containing 6.25μM folinic acid and 200μM of cysteine as a sulfur source. Transformants were grown in minimal medium containing methionine and folinic acid, harvested, washed and resuspended in water and serially diluted to give OD_600 nm_ values equal to 0.2, 0.02, 0.002 and 0.0002. A 10 μl aliquot of these dilutions were spotted on to minimal medium. The photographs were taken after 2 days of incubation at 30◦C.

Several important observations can be made from these results. Firstly, as one might expect, the same TMDs which are prominent in glutathione transport seem also to be important for folate uptake. And those TMDs, like TMD13 which were not observed to have any specific role glutathione uptake was also not important for folate uptake. Mutants defective in folate uptake and GSH uptake or both were re-evaluated using the previous described plate-based dual complementation-cum-toxicity assay (Supplementary Fig. S7) (Kaur et al., 2009; Kaur and Bachhawat, 2009). Secondly, most of the residues which were defective in glutathione uptake, were also observed to be defective in folate uptake. These residues may include those that are probably recognizing common features in the two substrates. It may also include residues involved in the translocation process, rather than in substrate recognition. Thirdly, we could observe residues whose mutation to alanine led to defect in glutathione transport, but not in folate transport. These include V185A in TMD3, G230A in TMD4, Y374A in TMD6 and Q526A in TMD9. Conversely, we observed residues whose mutations led to defect in folate transport but not in glutathione transport. These include residue L231A in TMD4, and the residues S427A, Y428A and L434A in TMD7. The conservation of these residues in orthologs could therefore be used as potential signatures in the search for folate transporters amongst the numerous OPT members.

### Phylogenetic analysis of the Oligopeptide Transporter (OPT) family

Opt1p belongs to the Oligopeptide Transporter (OPT) family. The OPT family is divided into the PT clade and the YSL clade. Opt1p belongs to the PT clade (or Peptide Transporter clade) and these members are found only in yeasts, fungi and plants (Fig.5). The function of a few of these have been identified in glutathione transport and in oligopeptide transport. However, the majority of these members are orphan transporters. The YSL (Yellow Striped-Like) clade is a more remote set of transporters that are found not only in plants, yeast and fungi but also in bacteria. Many of the members of the YSL clade function as transporters of metal chelating peptides and metal-nicotinamine complexes. *S. cerevisiae* has two members of the PT family, Opt1p and Opt2p, out of which Opt2p seems to be primarily localized to golgi with a role in lipid bilayer asymmetry (Yamauchi et al., 2014). Most other yeasts have multiple members in the PT clade. For example, *C. albicans* and *A. fumigatus* have 8 OPT’s each. We initially attempted to use the residues solely essential for folate uptake in Opt1p as signatures that could potentially identify folate transporters among the OPT members. However, as no clear signature was immediately apparent in the different organisms-specific alignments (Supplementary Fig. S8), we decided to evaluate OPT members from different organisms to examine if they might be folate transporters.

**Fig.5:**
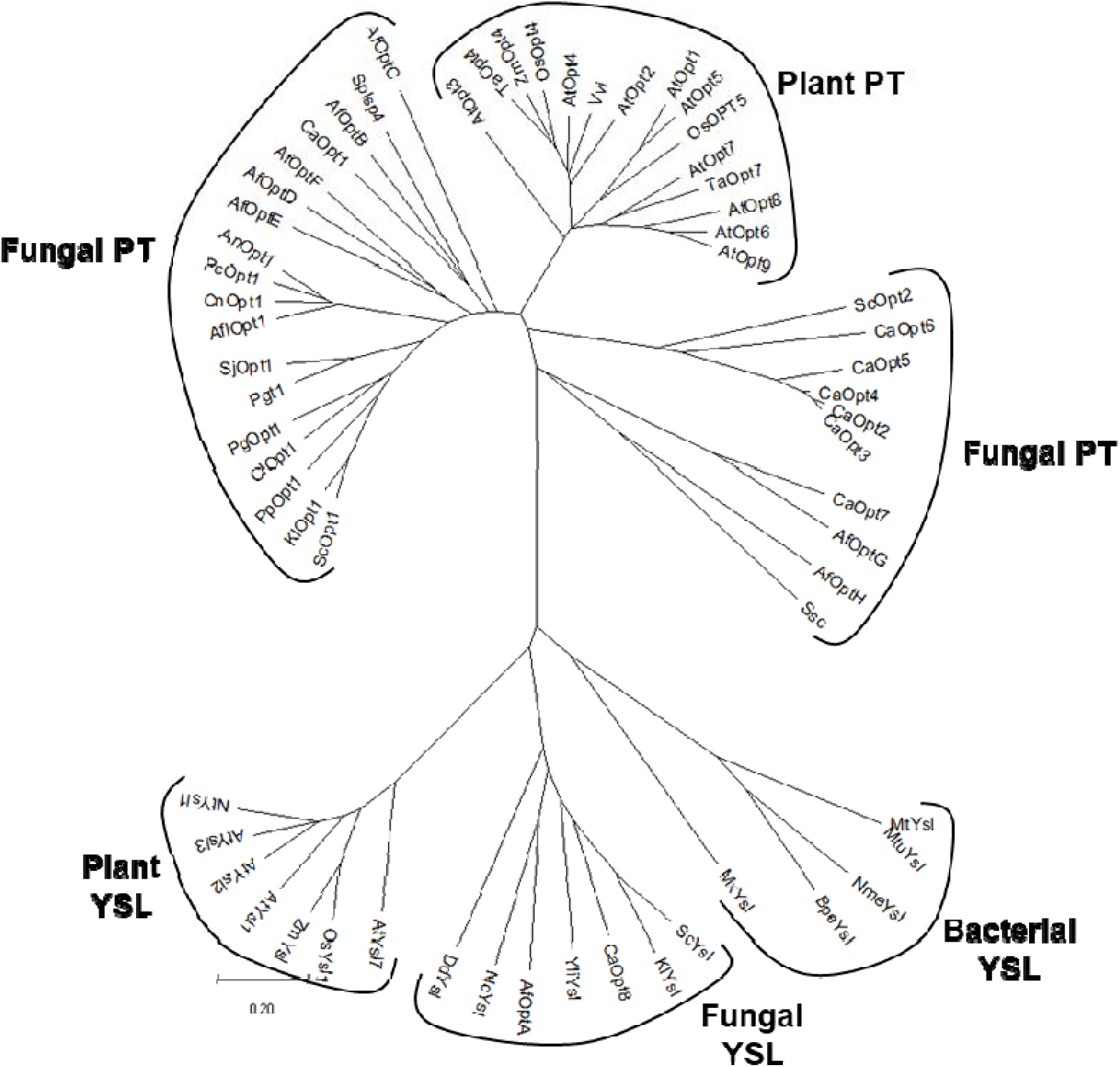
Unrooted phylogenetic tree of the OPT family. Phylogenetic tree of the representative members (Supplementary table S4) from the OPT family nd includes members of both the PT clade and the YSL clade. The tree was drawn with MEGA 11 software.

### *Candida albicans CaOPT1* and *Aspergillus fumigatus* OPTB and OPTH transport folate

*Candida albicans* is a fungal opportunistic pathogen which cause systemic fungal infections in humans (Hazen, 1995). *C. albicans* has eight oligopeptide transporter with seven of them, CaOPT1–CaOPT7, belonging to the peptide transporter (PT) clade. The transporters are named CaOPT1 to CaOPT8 in decreasing order of similarity to CaOpt1p, the first oligopeptide transporter identified in *C. albicans* (Reuß and Morschhäuser, 2006). CaOpt1p, is the apparent orthologue to ScOpt1p of *S. cerevisae*, sharing 39% of sequence similarity with ScOpt1p. Despite this similarity, *CaOPT1* was unable to transport GSH in *C. albicans* (Desai et al., 2011). Instead, a more distant homolog, CaOpt7p, with only 25% sequence identity, functions as the glutathione transporter, as previously demonstrated through uptake assays (Desai et al., 2011).

Based on the higher sequence identity of CaOPT1 with ScOPT1 and yet the functional similarity of CaOPT7 with ScOPT1, both CaOPT1 and CaOPT7 were tested for their ability to transport folate using the growth assay. CaOPT1 and CaOPT7 expressed under a TEF promoter were transformed in a *met15*Δ *fol2*Δ strain and assessed for growth on SD medium containing folinic acid along with different sulfur sources. It was observed that CaOPT1 but not CaOPT7 supported growth on folate. Interestingly, the growth on folinic acid with the CaOPT1 transporter was not inhibited by glutathione even at 200μM (Fig.6A), in contrast to ScOPT1. We further confirmed the ability of *C. albicans* CaOPT1 to transport folate by uptake assays (Fig.6C) which indicates that CaOPT1 is indeed a folate transporter.

**Fig.6:**
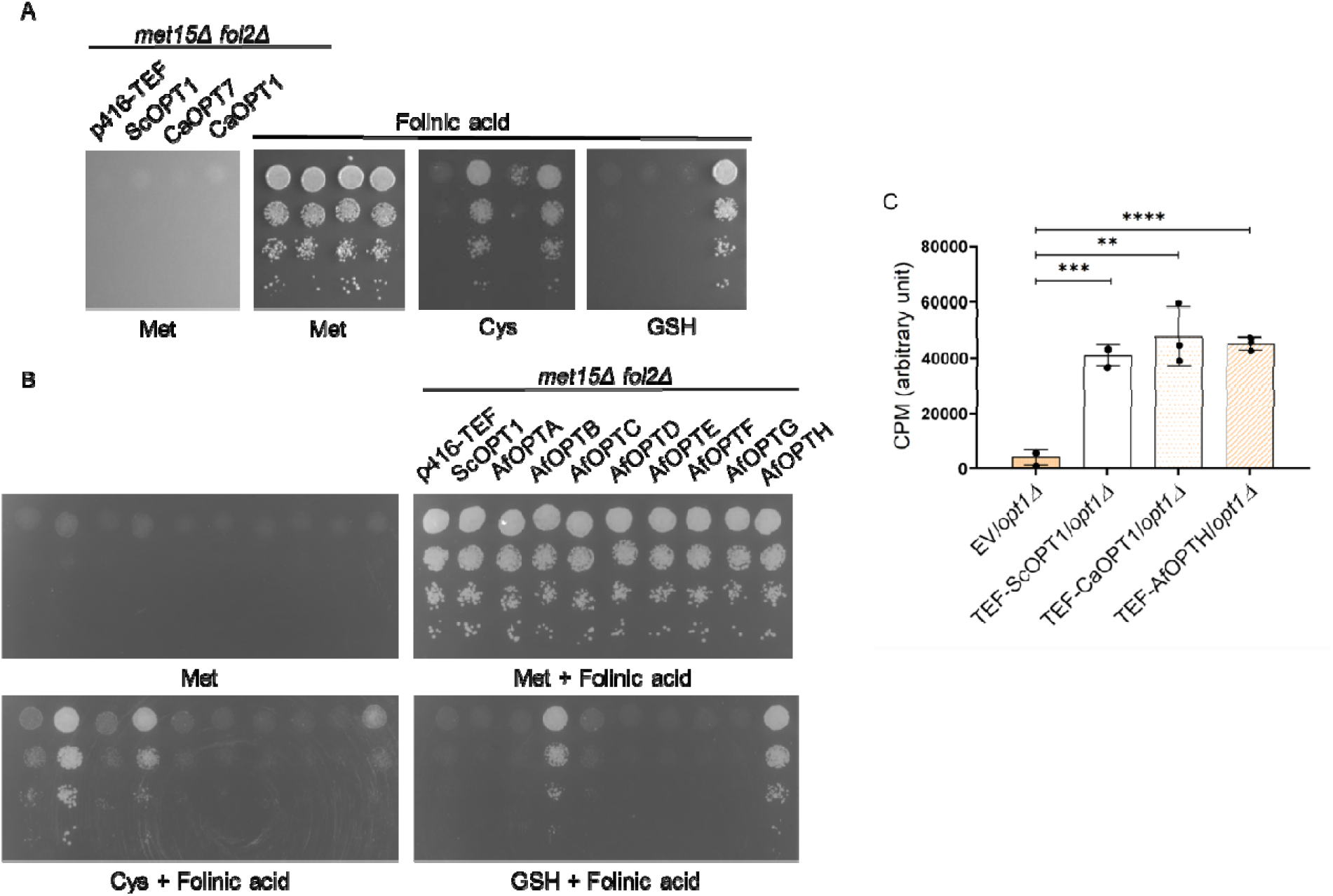
*C. albicans* and *A. fumigatus* OPT members are folate transporter. A. Functional complementation of CaOPT1 and CaOPT7 was performed using the growth assay to evaluate its role in folinic acid transport. Plasmids carrying CaOPT1 under the TEF promoter, along with the corresponding empty vectors, were transformed into the *S. cerevisiae met15*Δ *fol2*Δ strain. Dilution spotting assays was performed at 100μM folinic acid concentrations using 200μM of different sulfur sources. Photographs were taken after two days of incubation at 30 °C. B. Functional complementation of AfOPT clones were performed using yeast growth assay to evaluate its role in folinic acid transport. Plasmids carrying AfOPT1 under the MET25 promoter, were transformed into the *S. cerevisiae met15*Δ *fol2*Δ strain. Dilution spotting assays were performed on medium 50μM or 100μM folinic acid along with 200μM of different sulfur sources. Photographs were taken after two days of incubation at 30 °C. C. Intracellular folinic acid levels were quantified using LC-MS/MS in the *opt1*Δ strain with vector control and CaOPT1 and AfOPTH overexpression after 5 minutes of supplementation with 100 µM of folinic acid.

*Aspergillus fumigatus* is a fungal pathogen that has eight OPT members, named AfOPTA-AfOPTH, which have been otherwise annotated as putative peptide transporters (Hartmann et al., 2011). All eight AfOPT proteins were assessed for their role in folate transport using the growth assay. The AfOPTs that we obtained were under the control of the methionine-repressible MET25 promoter (Hartmann et al., 2011). These were transformed into the *S. cerevisiae met15*Δ *fol2*Δ strain and assessed using the growth assay. AfOPTB and AfOPTH were observed to confer growth on folinic acid containing media indicating their potential role as folate transporters (Fig.6B). Additionally, the presence of glutathione (GSH) did not inhibit the growth of AfOPTB and AfOPTH when utilizing folinic acid, suggesting that glutathione does not compete with folinic acid in these transporters, similar to what was observed with CaOPT1. In fact, none of the *A. fumigatus* OPTs appeared to transport GSH based on the lack of complementation by AfOPT clones in the *met15*Δ *opt1*Δ strain on GSH-containing media (Supplementary Fig. S9) (Kaur and Bachhawat, 2009; Thakur and Bachhawat, 2010). AfOPTH showed better rescue and we therefore carried out uptake experiments with AfOPTH to confirm its ability to transport folates (Fig.6C).

### OPT homologs of *Arabidopsis thaliana* can transport folates

*A. thaliana* has nine OPT members within the PT clade of the OPT family (Koh et al., 2002), of which AtOPT4 has been previously been shown to utilize GSH as a sulfur source when present at 400 uM concentrations. GSH transporter (Zhang et al., 2016). To investigate whether OPT1 homologs in *A. thaliana* can transport folate, they were cloned into yeast expression vectors and evaluated using the growth assay. cDNA of the *A. thaliana* OPTs from, AtOPT1 to AtOPT8 were procured from Arabidopsis Biological Resource Center (ARBC) and cloned under the TEF promoter. (Full length AtOPT9 clone was not available in ARBC and therefore not included in the current study). These were introduced into the *met15*Δ *fol2*Δ strain and assayed using the growth assay.

It was observed that AtOPT2 and AtOPT4 supported growth on folinic acid-supplemented medium. In the case of AtOPT6, functional complementation was observed only after extended incubation (Supplementary Fig. S10) and exclusively when glutathione served as the sulfur source. To further investigate the role of OPT6 in folate transport, we evaluated AtOPT6 (expressed using a multic py plasmid), which resulted in some growth relative to the vector control. However, this growth remained restricted to conditions where glutathione was used as the sulfur source (Fig.7A).

**Fig.7:**
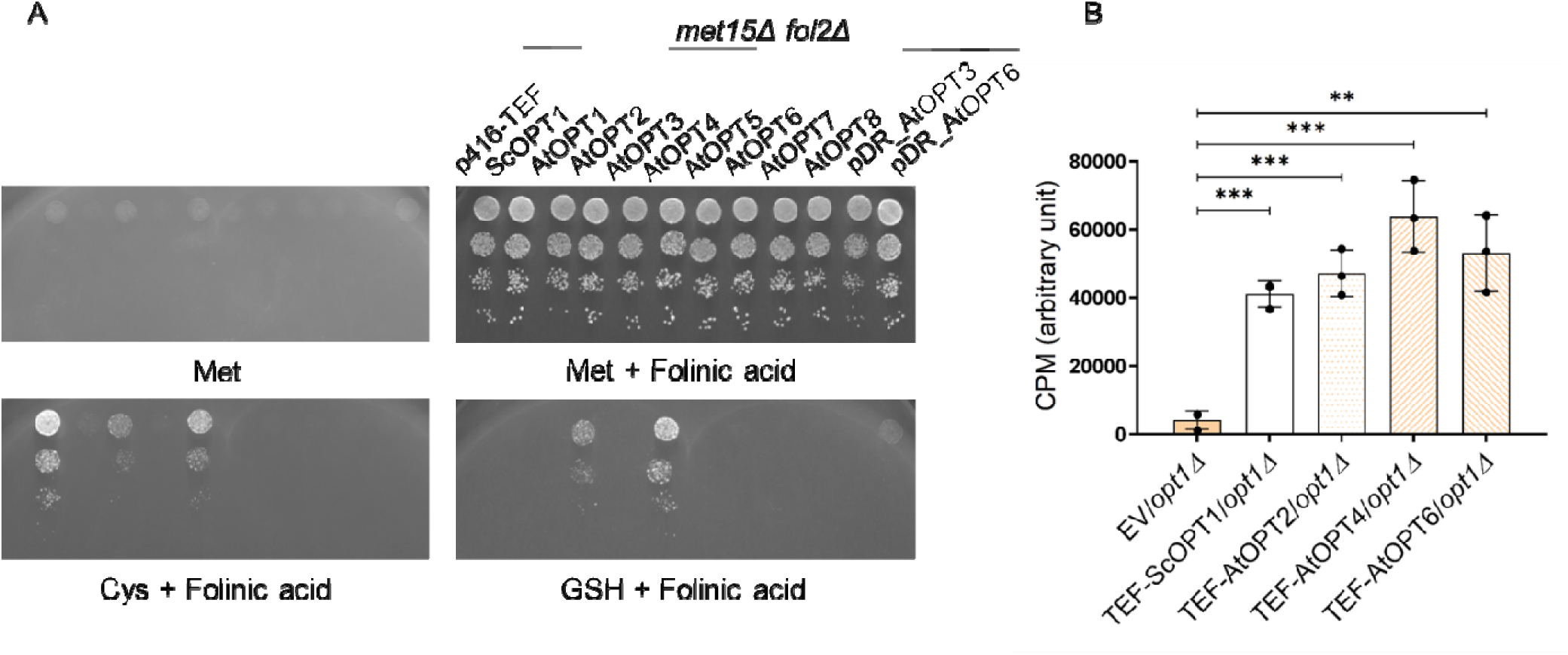
*Arabidopsis thaliana* AtOPT2, AtOPT4 and AtOPT6 are folate transporters. A. Plasmids carrying AtOPTs under the TEF promoter, were transformed into the *S. cerevisiae met15*Δ *fol2*Δ strain and dilution spotting assays was performed in 6.25μM of folinic acid media with 200μM of different sulfur sources. Photographs were taken after two days of incubation at 30 °C. B. Intracellular folinic acid levels were quantified using LC-MS/MS in the *opt1*Δ strain with vector control and AtOPT overexpression after 5 minutes of supplementation with 100µM of folinic acid.

To evaluate folate uptake by AtOPT2, AtOPT4, and AtOPT6, the clones (downstream of TEF) were transformed into *opt1*Δ. Folinic acid uptake was assessed by quantifying intracellular levels following a five-minute incubation with 100 µM folinic acid (Fig.7B). The results indicate that AtOPT2 and AtOPT4 and even AtOPT6 contribute to folate transport.

### Folate transporters of fungi and plants efficiently utilize MTHF

The most abundant folate in nature is methyl tetrahydrofolate. This has been observed in yeasts (Gmelch et al., 2020), humans (Pfeiffer et al., 2004) and in plants (Fyfe et al., 2022) and in all cases the major fraction of all the various folate vitamers, MTHF was the most abundant. It was therefore important to evaluate how the pathogenic yeast and fungi can utilize this folate. To evaluate this, the various folinic acid transporting homologs of yeast and fungi and *A. thaliana* were evaluated on both folic acid and methyl tetrahydrofolate.

We find that most of the folate transporters supported significant growth with MTHF (Fig 8). *C. albicans* CaOPT1 and *A. thaliana* AtOPT4 showed significantly higher growth than even ScOPT1. *A. fumigatus* OPTH also grew well on MHTF comparably to ScOPT1 and *Kluvyeromyces lactis*, KlOPT1. Only AtOPT2, AtOPT6 and AfOPTB showed negligible growth on methyl tetrahydrofolate (Fig.8). This indicates that many of these transporters have the capability of transporting folates and MTHF, and given the wide availability of MTHF in natural environments, the importance of these transporters in their natural environments becomes physiologically relevant.

**Fig.8.**
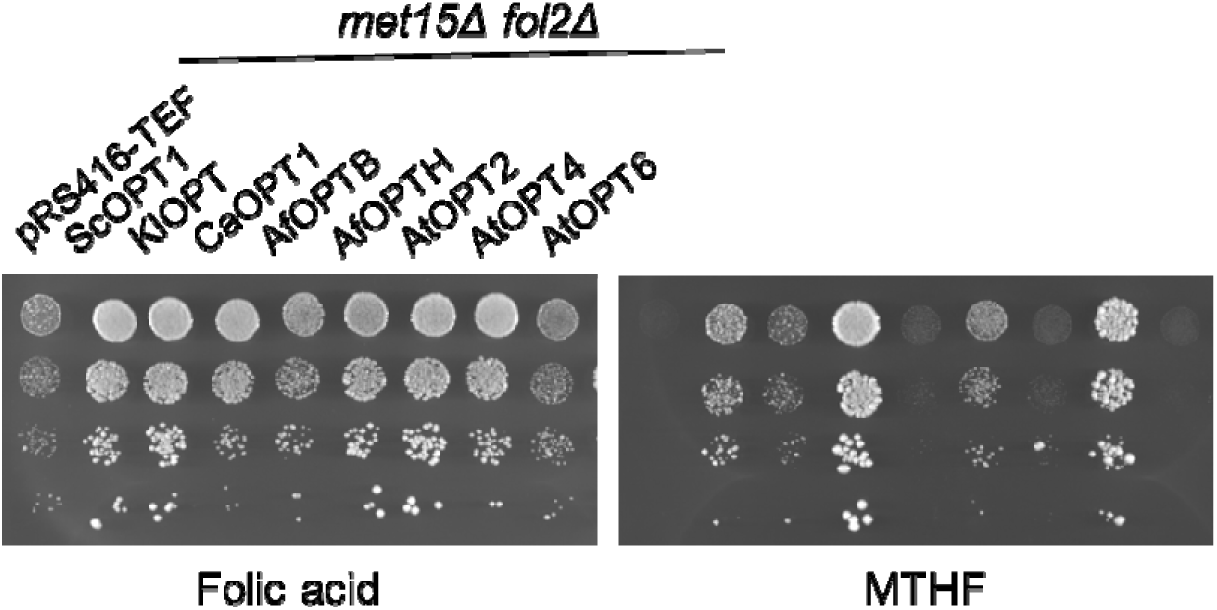
Ability of OPT homologs to grow on different forms of folate i.e. folic acid and MTHF. Plasmids carrying OPT homologs under the TEF promoter, were transformed into the *S. cerevisiae met15*Δ *fol2*Δ strain and dilution spotting assays was performed on 100μM of each folic acid and MTHF media with 200μM of methionine as a sulfur sources. Photographs were taken after two days of incubation at 30 °C.

## Discussion

Plasma membrane transporters of folate have not been identified in yeast, fungi and plants, and it was also speculated that a plasma membrane transporter for folate probably may not exist (Güldener et al., 2004). In this study we identify folate transporters and show that many members of the oligopeptide transporter (OPT) family (the PT clade of this family is unique to these organisms) include folate transporters. All these organisms each have a large number of OPT members. Hoewever, only a small sub-set of the fungal transporters has been shown to be involved in high affinity glutathione transport. Most members of the oligopeptide transporter family have no assigned function, and our discovery that many of them are capable of transporting folate efficiently, assigns a critical function to some of these orphan transporters.

*S. cerevisiae* has the best characterized transporter of this family, ScOpt1 or Hgt1p. It has been extensively investigated as a high-affinity plasma membrane GSH transporter, although it also transports oligopeptides with low-affinity (Bourbouloux et al., 2000; Hauser et al., 2000). It was therefore surprising that this transporter could also transport folates. Both folates and glutathione are tripartite structure with a common glutamate moiety in both. However, beyond this comparison there appears no real similarity in the two structures since folates are made of the pterin moiety, the 4-amino benzoate moiety and glutamate, while glutathione is a tripeptide composed of glycine, cysteine and glutamate. Despite these apparently distinct structures, there are common recognizable features. When we examined the literature, we observed that the human multidrug resistance protein MRP3, an ABC transporter closely associated with the MDR family, that effluxes glutathione conjugates were also shown to efflux methotrexate (MTX), a folate analog, along with folic acid (FA) and N5-formyltetrahydrofolic acid (leucovorin). This was in addition to the efflux of glutathione and glucuronate conjugates. Most interestingly, MRP3 was found to have a higher affinity for the folate forms than for GSH and its conjugates (Zeng et al., 2001, 2000, 1999). Another vacuolar MRP in plants, AtMRP1 identified in *A. thaliana*, has been shown to transport both folate and glutathione conjugates (Raichaudhuri et al., 2009).

How these transporters recognize these two structurally diverse substrates is intriguing. We investigated this in detail in Opt1p as we were interested in obtaining insights about the residues involved in specificity of recognition which could help to identify folate transporters in this large oligopeptide transporter family. We experimentally evaluated this using an alanine-scanned mutant library of all the TMDs in Optip (Zulkifli et al., 2016). Using this resource, we observed that many of the residues that were important for glutathione uptake were also important for folate uptake. However, we also found residues that were important exclusively for either folate uptake or glutathione uptake. Thus, residues L429, L434, Y449, L231 and Y428 were found exclusively important for folate uptake while the residues V185, G230, Y374 and Q526 residues were exclusively important for glutathione uptake.It is possible that as more folate transporters are identified amongst the OPT members, more definitive signatures for these transporters might emerge based on these observations. It is interesting, in this context, that of the 5 residues (V185A/ TMD3, G230A / TMD4, Y374A / TMD6, Y449A / TMD7; Q526A /TMD9) shown to be specific for glutathione transport (but not folate transport), 4 of these V185, Y374, L429 and Q526 were in fact show to have a higher Km when these residues were mutated to alanine (Zulkifli et al., 2016). V185A had a 3-fold higher Km, Y374A had a 2-fold higher Km, L429A had a 4-fold higher Km and Q526A had greater than 10-fold higher Km as compared to the WT molecule, clearly implicating them in substrate recognition, and this is borne out by their specificity to glutathione but not to folate, validating the genetic data.

Methotrexate, the well-known antifolate, has been explored in many previous studies in yeast (Delitheos et al., 1995; Scholz and Jaenicke, 1968; Wintersberger and Hirsch, 1973) that includes a more recent genome-wide search for methotrexate resistant mutants (Wong et al., 2017). However, none of these studies led to the identification of OPT1 mutants, although our studies here show that Opt1 overexpression leads to enhanced methotrexate sensitivity. One possibility for the failure of previous studies to identify OPT1 using methotrexate might be due to the use of YPD medium, which contains glutathione (from yeast extract), and it could interfere with the transport of methotrexate as it does with folates in *S.cerevisiae*. It may also explain the higher resistances seen in these studies.

It is remarkable that the *S. cerevisiae* ScOPT1/Hgt1p is not only the major transporter of folates in this yeast, but also the sole transporter of glutathione. In fact, it has been previously shown that deleting glutathione biosynthesis along with the OPT1 transporter in *S. cerevisiae* led to inviable cells, and *gsh1*Δ *opt1*Δ was synthetically lethal (Bourbouloux et al., 2000), just as we observed that *fol2*Δ *opt1*Δ was also synthetically very sick in this study. Thus, the same transporter plays a seminal role in both glutathione and folate, two essential molecules of the cell.

Folate biosynthesis occurs in fungi and yeast, but not in humans. Thus, the folate pathway of these organisms has been proposed as a potential antifungal target (DeJarnette, 2020; Meir and Osherov, 2018). However, the demonstration here that *C. albicans* and *A. fumigatus* both have dedicated folate transporters raises the question that if fungi are able to source intracellular sources of folates using the transporters described then the efficacy of folate biosynthetic pathways as targets would be diminished. The most predominant folates in humans is methyltetrahdyrofolate which forms upto 93% of the total pool of folates (Pfeiffer et al., 2004). It is in interesting that the fungal pathogens *C. albicans* (CaOPT1) and *A.fumigatus* (OPTH) both have an ability to use these dominant folates, and CaOPT1 seemed to utilize MTHF much more efficiently than the *S. cerevisae* homolog. Future studies therefore can see if these transporters affect the pathogenesis of these fungi *in vivo*, and if they do, the antifungals would have to target not only the folate pathways but also the transporters.

In the case of *C. albicans*, we could observe an evolutionary diversification of function with respect to the oligopeptide transporter family. Thus CaOPT1 (the closest homologue to ScOPT1) to be a folate transporter, with no apparent ability of glutathione transport, while CaOPT7 a transporter of glutathione as shown earlier (Desai et al., 2011) but no ability to transport folate. We have not explored the roles of the other CaOPTs and it would be interesting to see whether any of them can also transport folates. It was interesting and also surprising to note that in *A. fumigatus*, two of the transporters could transport folate. Why two OPTs are required for folate transport in this fungus is not clear.

The plant folate biosynthetic pathway is compartmentalized across the cytosol, mitochondria, and chloroplasts, and folates are found throughout these cellular compartments (Ravanel et al., 2004, 2001). Therefore, in plants, in addition to cellular folate uptake, intracellular transport of folate or its precursors is also required (Hanson and Gregory, 2011). Three intracellular folate transporters have been characterized in plants—two in the chloroplast and one in the vacuolar membrane. While evidence suggests the presence of plasma membrane folate transporters in *A. thaliana* (Ishikawa et al., 2003; Prabhu et al., 1998), they have not been identified prior to this study. The findings reported here expands the understanding of folate metabolism and transport in plants. AtOPT2 and AtOPT4 interestingly, are phylogenetically closely related. These OPTs are expressed in the plasma membrane but each in different stages and structure of the plants. Folates are essential components of the human diet, and yet there is an immense worldwide problem of folate deficiency in pregnant women leading to birth defects and anemia. There are several attempts at folate biofortification of foods to alleviate this malnutrition, but the problem is complex (Blancquaert et al., 2014) and the most promising approach seems to be through metabolic engineering of plants (Storozhenko et al., 2007). However, the regulation of the two branches leading up to folate is complex, leading to varied results in different plants. In this situation, the folate transporters identified here could be a useful tool that could be exploited in some plants for developing folate enriched foods.

## Methods and Materials

Chemicals and Reagents-All chemicals used in the present study were of either analytical or molecular biology grades and were obtained from commercial sources. Media components were purchased from Difco, Merck, and HiMedia. Vitamin free YNB was obtained from HiMedia. Restriction enzymes, T4 DNA ligase and other DNA modifying enzymes were obtained from New England Biolabs. Phusion™ High-Fidelity DNA Polymerase was obtained from Thermo Scientific while Taq Polymerase was obtained from Promega. Gel extraction kits and plasmid miniprep columns were obtained from Bioneer Inc. (Daejeon, South Korea) or Agilent Technologies or Promega. Oligonucleotides were purchased from Merck and IDT. Folinic acid used in the study was of BioXtra grade and was obtained from Merk. Folic acid was obtained from Schircks Laboratories. Calcium L-5 methyl tetrahydrofolate was obtained from Anthem bioSciences. Methotrexate and glutathione were purchased from Merck.

Strains and Growth-*Escherichia coli* DH5α was used as cloning host. The list of *Saccharomyces cerevisiae* strains used in this study and their genotypes is shown in Supplementary Table 5. The yeast strains were maintained in YPD (yeast extract, peptone, and dextrose). *S. cerevisiae* W303 strains were maintained in YPAD (yeast extract, peptone, dextrose and adenine). 100 μM of folinic acid was used for maintaining *fol2*Δ strains.

For growth assays, synthetic defined (SD) minimal media containing yeast nitrogen base, ammonium sulfate, and dextrose supplemented with the required amino acids and bases (80 mg/l) were used or Synthetic defined (SD) minimal media containing vitamin free yeast nitrogen base and dextrose supplemented with the required vitamins (lacking folic acid), amino acids and bases (80 mg/l) was used. Minimal medium containing potassium hydrogen phthalate, sodium hydrogen phosphate, ammonium chloride with salts, vitamins (lacking folic acid), minerals and glucose was also used (Alfa, 1993).

Folinic acid and calcium L-5 methyl tetrahydrofolate was used when required at a concentration of 100μM, unless specified otherwise. Glutathione was added wherever necessary at required concentrations. Sporulation plates were prepared as described by (Kaiser et al., 1994) Growth, handling of bacteria and yeast, and all the molecular techniques used in the study were according to the standard protocols (Sambrook et al., 1989)

Yeast-based growth assay to screen for folate uptake-When OPT1 was overexpressed using the TEF vector and grown on SD medium with low levels of cysteine and folinic acid, the cells expressing OPT1 exhibited improved growth compared to the vector control. The YNB used for preparing the SD medium contained 2 μM folinic acid, which is not sufficient to support the growth of fol2Δ cells. However, on plates supplemented with both methionine and folinic acid, the growth of the vector control was comparable to that of the OPT1-expressing strain, likely because methionine can compensate for folinic acid auxotrophy. A similar growth advantage of OPT1 over the vector control was also observed in a complete synthetic medium with low concentrations of methionine and folinic acid. Nevertheless, SD medium was preferred over the complete synthetic medium due to its simplicity and consistent preparation. This observed growth difference was utilized to develop a yeast-based growth assay, in which OPT1mutants or homologs from other yeasts, fungi, and plants were heterologously expressed and screened.

Generation of FOL2 disruptions in different deletion strains-The FOL2 disruption cassette was generated as follows. Firstly, FOL2 gene (732bp) was cloned along with 321 bp flank in the 5’ end and 100bp flank at the 3’ end. The fragment was PCR amplified using the primer pair FP_Fol2_OH_BamHI and RP_Fol2_OH_XhoI using chromosomal DNA of a wild type strain (BY4741) as a temple. Primers used are listed in the Supplimentary table S7. The 1.2-kilobase PCR product obtained was digested with BamHI and XhoI and cloned into a single copy, URA3-based yeast expression vector downstream of the TEF promoter. The yeast HIS3 gene was PCR amplified using primer pair FP_fol2cas_EcoRI and RP_Fol2cas_EcoRI using pRS313 vector (ABE 3569, laboratory stock) as a template and cloned into the FOL2 gene using the single EcoRI site in the FOL2 gene. To generate linearized *fol2*Δ*::HIS3* disruption cassette, the above generated plasmid was digested using BamHI and XhoI. This FOL2 disruption cassette, contained the sequence of Histidine auxotrophic marker flanked at both sides with the FOL2 sequence of 657 and 496 at the 5’ and 3’ termini. The disruption cassette was transformed into the yeast cells and selected by histidine prototrophy. The disruptants were further confirmed by folinic acid auxotrophy.

Creation of an OPT1 deletion-The OPT1 gene was disrupted in the W303 MATa strain background using the one-step PCR-mediated gene disruption. The *opt1*Δ::LEU2 disruption cassette was generated using the primer pair FP_opt_delcas_Leu and RP_opt_delcas_Leu using plasmid pRS315TEF (ABE 3488, laboratory stock) as a template. The 1.7-kb PCR product obtained was transformed into W303 strains, and the transformants were selected on minimal medium without leucine. The transformants in *opt1*Δ strain were confirmed for the disruption by diagnostic PCR using the primer pair FP_opt_delcas_ck and RP_opt1_delcas_ck

Cloning of the *Arabidopsis thaliana* AtOPTs into yeast expression vector-*In-vivo* homology-based cloning was used to clone Arabidopsis thaliana OPT’s into the yeast expression vector pRS416TEF. The split-marker mediated multiple-piece cloning method was employed (Kuijpers et al., 2013) to increase the efficiency of fragment assembly and reduce chances of false positive (Singh et al., 2024). The vector pRS416_TEF (ABE443) (5461 bp), was split into 2 fragments such that one fragment contains the URA3 marker and bacterial origin of replication (F1 ori) (2267 bp) and the other fragment contained the CEN/ ARS region along with the ampicillin resistance gene (3484 bp). The two fragments were generated using primer pair FP_416URA and RP_416URA and FP_416CEN and RP_416CEN using pRS416_TEF as a temple. These chassis primers can be used universally, irrespective of gene(s) of interest. The junction primers, FP_416URA and RP_416CEN, were designed such that there is a homology of 284 bp between the URA fragment and the CEN fragment. A pair of primers were designed for PCR amplification of the At_OPT(1-8) so that they have homology with MCS region at their 5’ and 3’ ends. The forward primer, FP_AtOPT1-8_Frag, contains 34 bp of nucleotide sequence of TEF promoter followed by a 6 bp of restriction site sequence of any desired restriction endonuclease enzyme (for making the plasmid feasible for subcloning) which is further followed by a 20 bp of sequence from the gene of interest. The reverse primer RP_AtOPT1-8_Frag was also designed similarly. The AtOPT fragments was generated using the plasmid as a template ordered from Arabidopsis Biological Resource Center (ARBC). The DNA fragments were PCR amplified at the standardized annealing temperature conditions, and specific bands of respective lengths were eluted after gel-electrophoresis and were transformed along with the two vector fragments into the yeast cells. 200 fmol of DNA fragments were used for transformation into the yeast strain, BY4741 and were screened on synthetically defined (SD) media agar plates supplemented with amino acids lacking uracil.

Growth assays using dilution spotting-Growth assays using dilution spotting of *fol2*Δ (or WT) strains transformed with either Opt1 or its homologs or mutants cloned under pRS416 vector and its vector controls were transformed individually in *S. cerevisiae* BY4741 (sulfur auxotrophic) or BY4742. The transformants were selected and were grown overnight in a minimal medium with amino acid supplements and folinic acid as required. They were washed twice to remove the media components and reinoculated in a fresh medium containing amino acid supplements, without folinic acid, at an OD_600 nm_ of 0.20. The cells were folate-starved for 5–6 h until the early exponential phase was reached. These cells were harvested, washed, and resuspended in water to an OD_600 nm_ of 0.2. These suspensions were serially diluted to 1:10, 1:100, and 1:1000. 10μl were spotted on different minimal medium plates containing methionine/cysteine/ glutathione as sulfur sources and suboptimal folinic acid concentration (25μM). The plates were incubated at 30°C for 2–3 days, and then the images were taken using the Bio-Rad Gel Doc™ XR+ imaging system.

Yeast DNA Isolation and Yeast Transformation-Yeast chromosomal DNA was isolated by the glass bead lysis method and yeast transformations were carried out using the lithium acetate (Alfa, 1993; Amberg, 2005; Gietz and Schiestl, 1995)

Tetrad Analysis-To check the viability of *fol2*Δ and *opt1*Δ double deletes, a diploid heterozygous for both markers was constructed by crossing W303MATα (*fol2*Δ::HIS3) with W303 MATa (*opt1*Δ::LEU2). The diploid was sporulated on minimal sporulation plates and tetrads were dissected. The spores were dropped on YPAD medium containing 100uM folinic acid. After initial patching on YPAD plates supplemented with folinic acid, spores were replica plated on SD minus histidine and minus leucine plates. The disruption at the FOL2 locus was followed by folinic acid auxotrophy and histidine prototrophy, while disruption at OPT1 locus was followed by leucine prototrophy.

Transport Experiments-To measure uptake of folinic acid and calcium L-5-methyltetrahydrofolate in Saccharomyces cerevisiae, BY4741 transformants overexpressing ScOPT1 or its homologs under the TEF promoter were cultured overnight in a liquid minimal medium supplemented with amino acids but lacking folates. Cultures were incubated at 30 °C with shaking at 200 rpm for 12 hours. Cells were then reinoculated into fresh medium (again without folates) at an initial OD _nm_ of 0.20. After reaching the exponential growth phase, 100 μM of folinic acid or calcium L-5-methyltetrahydrofolate was added. The flasks were further incubated at 30 °C, shaken at 200 rpm, and samples were collected either at various time points for time-dependent uptake studies or after a single 5-minute incubation for endpoint analysis. Following incubation, metabolite quenching and extraction were performed as previously described (Walvekar et al., 2018). Briefly, 5.O.D. cells were added to ice-cold 60% methanol to halt metabolism, followed by centrifugation at 600g for 3 minutes at 4 °C. This was followed by another round of washing using 60% methanol. For extraction, 550μl of 75% (v/v) ethanol (pre-chilled to −45 °C) was added to the pellet, vortexed for 15 seconds, and incubated at −45 °C for 15 minutes. After another vortexing step (15 seconds), the samples were centrifuged at 21,000g for 10 minutes at −5 °C. The supernatant was collected into fresh tubes, recentrifuged under the same conditions, and then dried using a SpeedVac. The dried extracts were stored at −20 °C until analysis.

Quantification of folates was performed using LC-MS/MS as detailed by Walvekar et al. (2018), employing a Synergi™ 4-μm Fusion-RP 80-Å (100 × 4.6 mm) LC column and an AB Sciex QTRAP 6500 triple-quadrupole mass spectrometer.

Multiple sequence analysis and phylogenetic analysis-The OPT sequences were retrieved from Entrez. The multiple sequence alignment of the protein sequences was generated using the Clustal Omega (Sievers and Higgins, 2014), and a phylogenetic tree was constructed via the neighbor-joining method using MEGA 11 software (Tamura et al., 2021)

Statistical analysis-All uptake data has been analysed using student’s t test with p value cut off of 0.05 using Graph Pad Prism.

Modelling the yeast OPT1-Glutathione/Folinic acid complexes

The full structure of yeast OPT1 was modelled using the alphafold2 structure (loops) and the model (The transmembrane helices) generated and validated in our previous study (Zulkifli et al., 2016) using Modeller10.5. Loop refinement module was used in slow refinement mode to optimize the regions where the helices and the loops meet. The structures were subjected to structure quality assessment programs Procheck (Laskowski et al., 1993), Verify 3D (Bowie et al., 1991) and Errat (Colovos and Yeates, 1993) for validation. The yeast OPT11 model was then subjected to protein preparation using the Protein Preparation Wizard (PPW) module (Protein Preparation Wizard; Epik, Schrödinger, LLC, New York, NY, 2019) of Schrödinger software package, version 2019–2. Missing hydrogens were added and appropriate bond orders were assigned to the structures. The protonation states of the polar residues were optimized with the protassign module of PPW, which uses PROPKA to predict pKa values (pH 7.0 ± 2.0) and side chain functional group orientations. The structure was then subjected to restrained minimization (cutoff RMSD 0.3Å) with impref to avoid steric clashes. The prepared structure was used for preparation of grids and molecular docking. ‘Receptor Grid Generation’ module of Schrödinger was utilized to define an interaction grids for molecular docking. The interaction grid was defined in such a way that the whole α-helical transmembrane region was included in the box. Sizes of the inner and outer boxes were fixed to 16 Å and 22 Å respectively, keeping the centroid of the α-helices as the grid box centre. The structures of glutathione and folinic acid ware downloaded from PubChem database and prepared with LigPrep (LigPrep, Schrödinger, LLC,New York, NY, 2019), generating different ionization states at pH 7.0(± 2.0) using Epik ionizer and the best energy conformer was retained for each ligand for docking. The ligands were docked to yeast OPT1 structure (Grid) using the Glide module of Schrödinger software package with extra precision (XP) mode.20 OPLS_2005 force field was used for docking with all default parameters and the best docked pose for each ligand was retained for generation of the respective protein-ligand complexes.

## Acknowledgements

The work was supported by a grant-in-aid project from the Department of Biotechnology (BT/PR39040/BRB/10/1882/2020) to AKB and a DBT-Wellcome Trust India Alliance Senior Fellowship (IA/S/21/2/505922) to SL. NV was a recipient of a Senior Research Fellowship from the Council of Scientific and Industrial Research (CSIR), India, MS thanks SERB NPDF2022(PDF/2022/000700) for funding. We thank Sven Krappman for the *Aspergillus fumigatus* clones and Cecille Gaillard for the AtOPT6 and AtOPT3 clones on a multicopy plasmid. We thank NCBS/InStem/CCAMP Mass Spectrometry facility for liquid chromatography-tandem mass spectrometry used for the uptake studies.

