## Supplementary data for "Plasma membrane folate transport in fungi and plants is mediated by members of the oligopeptide transporter (OPT) family"

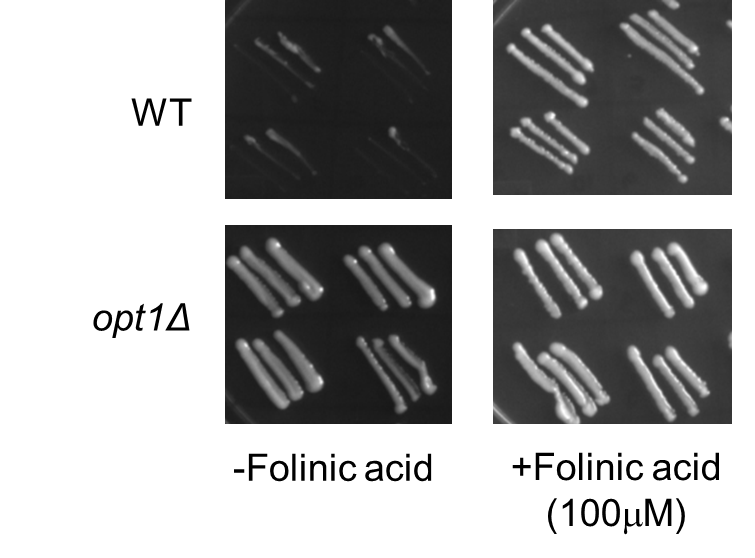

**Supplementary Fig. S1**: **The His^+^ transformants of *opt1Δ* strains (transformed with *fol2Δ*::HIS3 cassette) did not produce folinic acid auxotrophs**

His^+^ colonies appearing on WT and *opt1Δ* strains transformed with the *fol2::HIS3* disruption cassette, were streaked on vitamin free SD medium with and without folinic acid. The His^+^ transformants in *opt1Δ* grow with or without folinic acid, suggesting that FOL2 has not been disrupted.

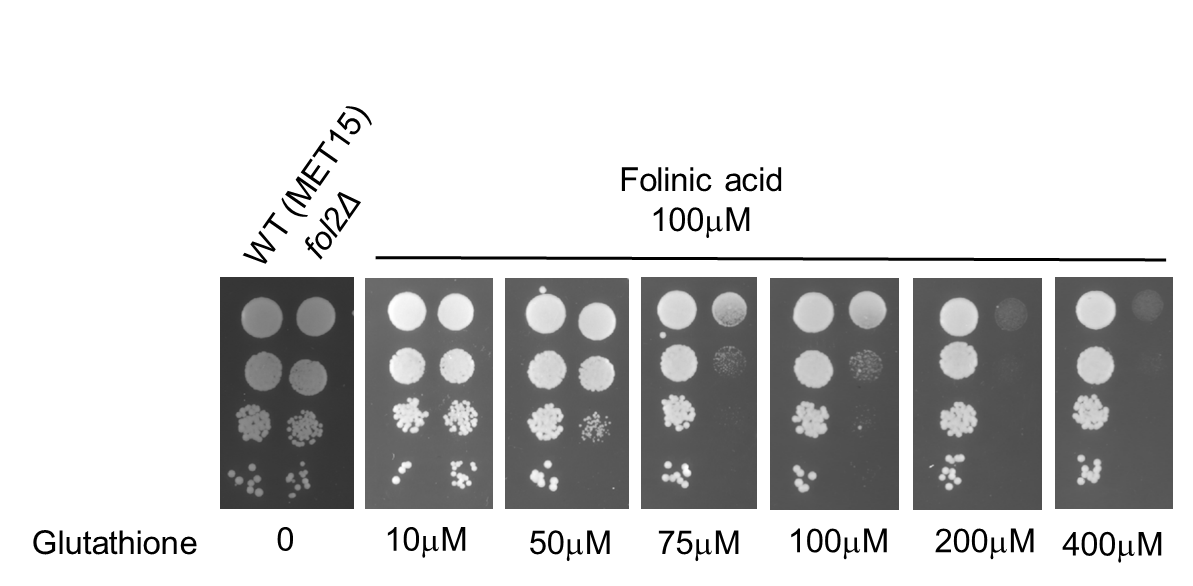

**Supplementary Fig. S2**: **Inhibition of the utilization of folates by GSH in the *fol2Δ* strain**

A dose-dependent inhibition of the *fol2Δ* strain by GSH was assessed in the *fol2Δ* strain (BY4742 background) on SD medium supplemented with 100μM folinic acid along with different concentrations of glutathione

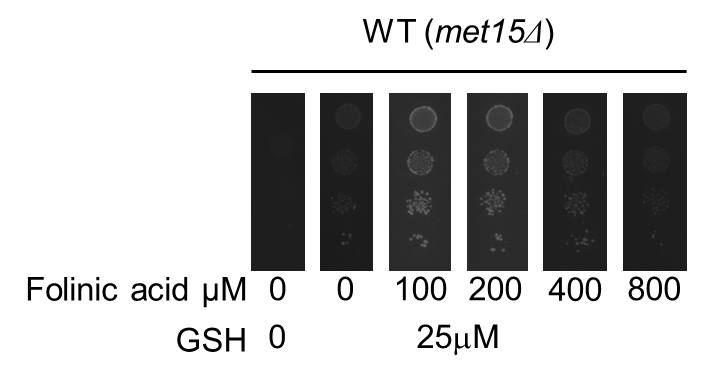

**Supplementary Fig. S3**: **Folinic acid inhibits glutathione utilization in the *met15Δ* strain**

The competition between GSH and folinic acid for uptake in the BY4741 strain was evaluated using a plate-based spot assay on SD medium supplemented with 25μM GSH and increasing concentrations of folinic acid

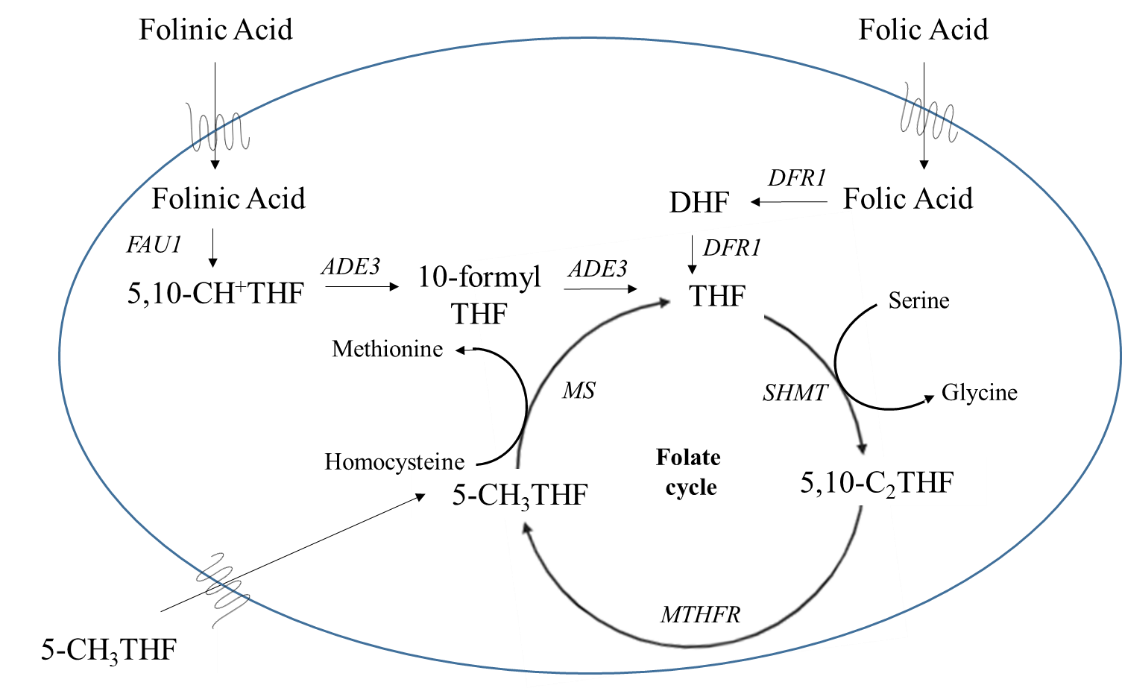

**Supplementary Fig. S4**: **Schematic representation of the uptake, utilization, and interconversion of various folate forms within the yeast cell**

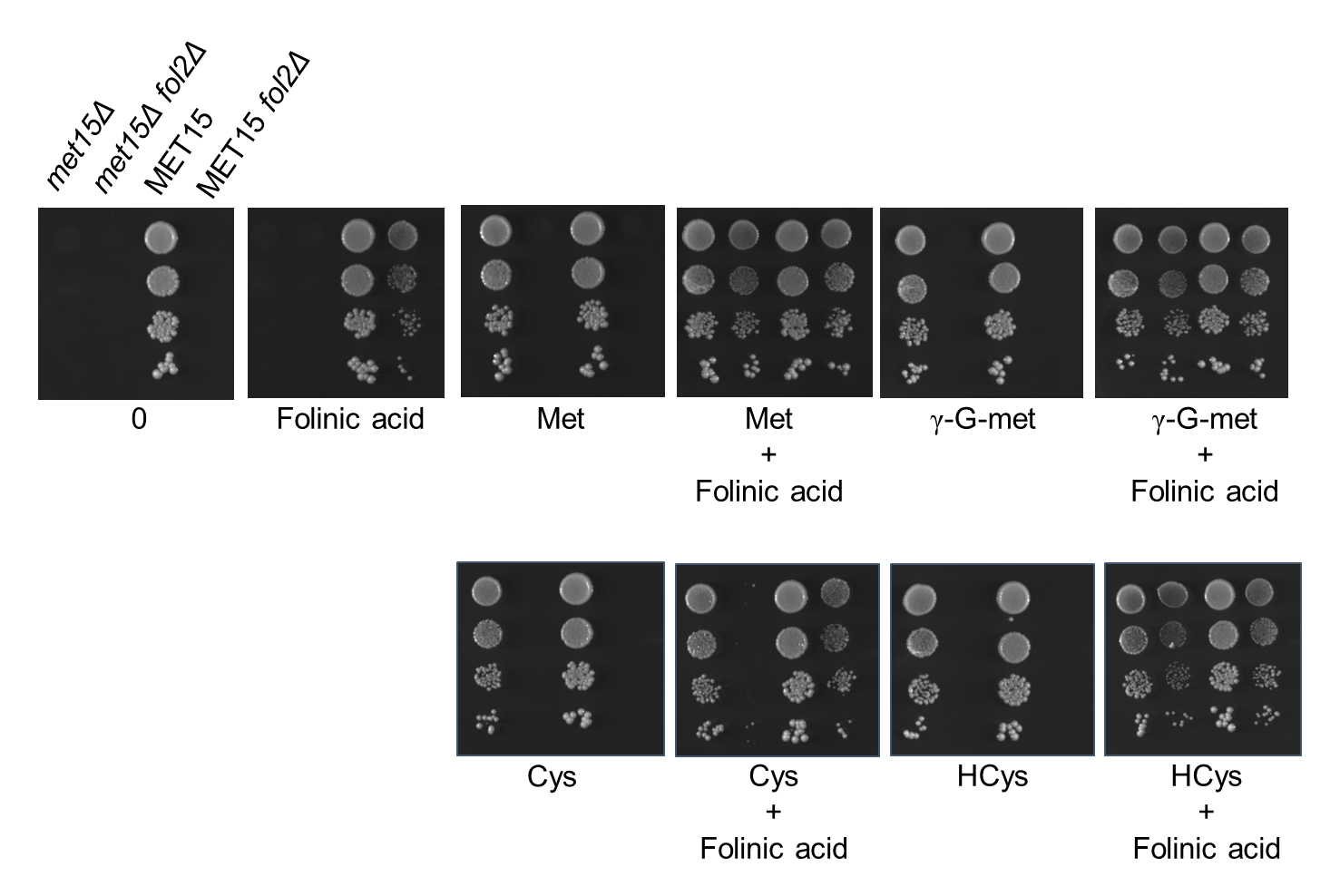

**Supplementary Fig. S5**: **Development of a growth assay to evaluate the role of OPT mutants and OPT members in folate transport**

Growth of *met15Δ* or MET15 strain along with their respective FOL2 deletions. Dilution spotting assays was performed with or without 100μM folinic acid using 200 μM of different sulfur sources. Photographs were taken after two days of incubation at 30 °C.

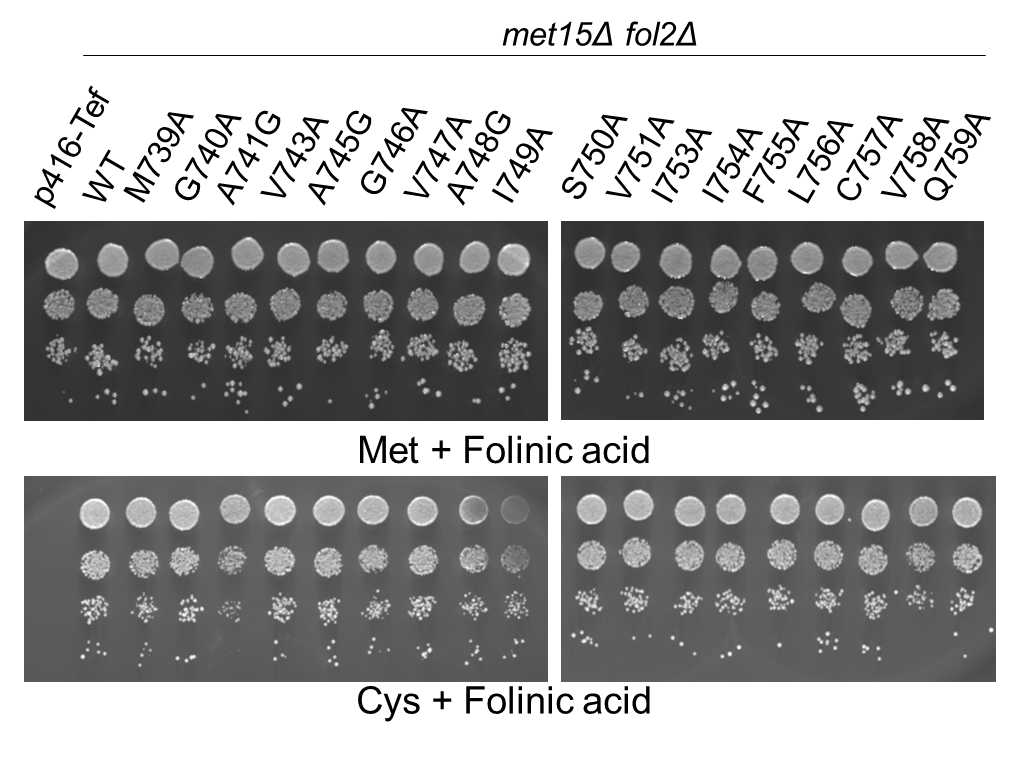

**Supplementary Fig. S6: TMD13 alanine mutants of Opt1p in folinic acid transport**.

The different alanine mutants of Opt1p under the TEF promoter and corresponding control vector (p416TEF) were transformed into strain *fol2Δ met15Δ* strain and evaluated for the ability to transport folinic acid using the growth assay by dilution spotting on minimal medium containing 6.25μM of folinic acid and 200 μM of cysteine as a sulfur source. Transformants were grown in minimal medium containing methionine and folinic acid, harvested, washed and resuspended in water and serially diluted to give OD_600 nm_ values equal to 0.2, 0.02, 0.002 and 0.0002. A 10 μl aliquot of these dilutions were spotted on to minimal medium. The photographs were taken after 2 days of incubation at 30◦C.

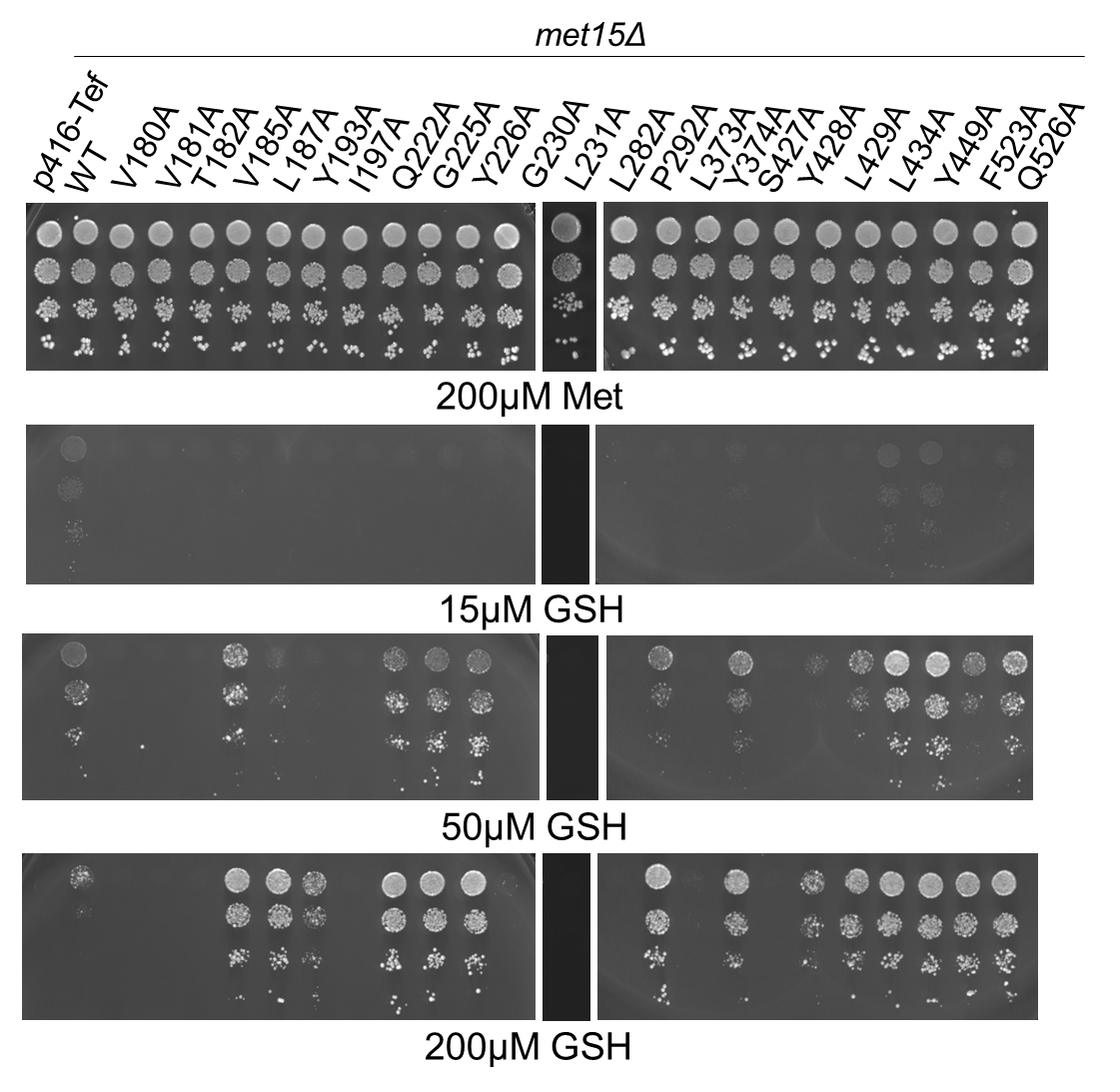

**Supplementary Fig. S7**: **Re-evaluation of Opt1p mutants impaired in folate or GSH transport for defects in glutathione utilization**

OPT1/HGT1 and the alanine mutants that showed defect in either folate or glutathione transport or both were reassessed for GSH utilization. The mutants along with vector control and WT protein were expressed under the TEF promoter were transformed in *met15Δ opt1Δ* strain. Transformants were subjected to plate-based dual complementation-cum-toxicity assay (Kaur, 2009) by serial dilution spotting on minimal medium containing different concentrations (15, 50 and 200 μM) of glutathione or 200 μM methionine (control). The photographs were taken after 2–3 days of incubation at 30◦C. The experiment was repeated with three independent transformations.

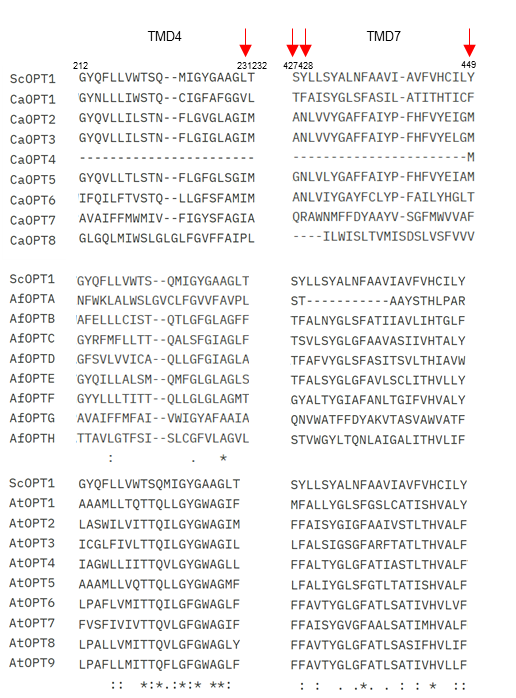

**Supplementary Fig. S8**: **Multiple sequence alignment of the protein sequences of the TMD4 and TMD7 regions of the PT members of *Candida albicans*, *Aspergillus fumigatus* and *Arabidopsis thaliana* OPT family**

Sequences were retrieved from Entrez at NCBI website and aligned using CLUSTALW program. The sequence alignment has been edited to show only the transmembrane domains 4 and 7. Residues corresponding to position 231, 427, 428, and 429 have been marked by an arrow.

**Yeast and fungi**: ScOPT1 (NP_012323.1), CaOPT1 (XP_718267), CaOPT2 (XP_713434), CaOPT3 (XP_713437), CaOPT4 (XP_710713), CaOPT5 (XP_717657), CaOPT6 (XP_722726), CaOPT7 (XP_713354), CaOPT8 (XP_718105), AfOPTA (XP_752732.1), AfOPTB (XP_755859.1), AfOPTC (XP_754401.1), AfOPTD (XP_753269.1), AfOPTE (XP_750894.1), AfOPTF (XP_750911.1), AfOPTG (XP_747762.1), AfOPTH (XP_746839.1)

**Plants**: AtOPT1 (NP_200404.1), AtOPT2 (NP_172464.1), AtOPT3 (NP_567493.5), AtOPT4 (NP_201246.1), AtOPT5 (NP_194389.1), AtOPT6 (NP_194503.1), AtOPT7 (NP_192815.1), AtOPT8 (NP_200164.1), AtOPT9 (NP_200163.1)

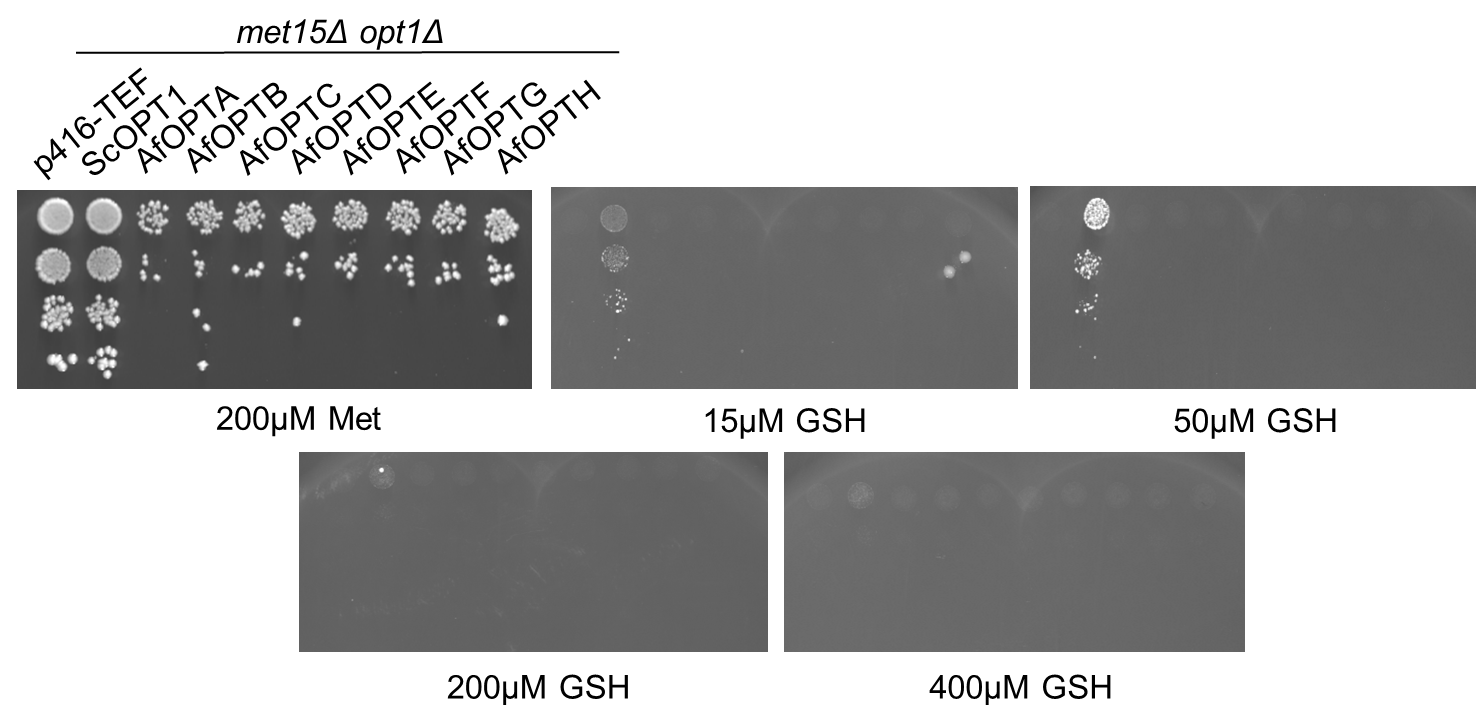

**Supplementary Fig. S9**: ***Aspergillus fumigatus OPT does not function as a GSH transporter***

Functional complementation of AfOPT was performed using a genetic screen to evaluate its role in GSH transport. Plasmids carrying AfOPT1 under the MET25 promoter, were expressed in a *met15Δ opt1Δ* background. Dilution spotting assays was performed at 15μM to 400μM GSH concentration. Photographs were taken after four days of incubation at 30 °C.

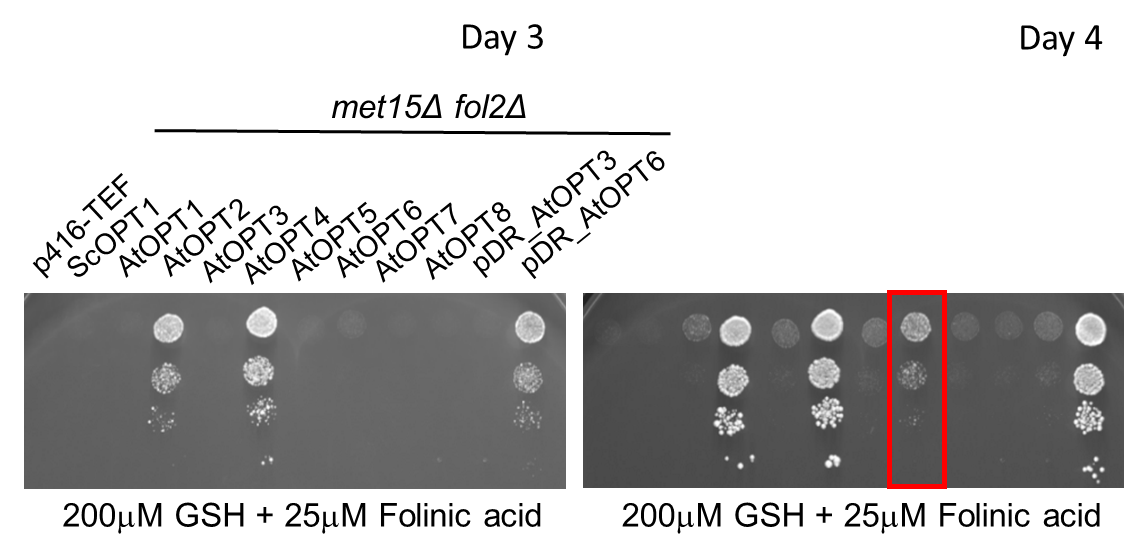

**Supplementary Fig. S10: *Arabidopsis thaliana* AtOPT6 is a folate transporter**

Plasmids carrying AtOPTs under the TEF promoter, were transformed into the S. cerevisiae *met15Δ fol2Δ* strain and dilution spotting assays was performed in 25 μM folinic acid media with glutathione as a sulfur sources. Photographs were taken after two days of incubation at 30 °C.

**Supplementary Table 1.** Summary of the total number of transformants and the number of transformants exhibiting folate auxotrophy in WT and *opt1Δ* strains when transformed with FOL2 disruption cassette

| Strain | No. of transformants | No. of tranformants assessed | No. of transformants showing folate auxotrophy | Fol2 disruption frequency% |
| --- | --- | --- | --- | --- |
| BY4741 (WT) | 1230 | 21 | 17 | ~85% |
| BY4741 *opt1Δ* | 68 | 68 | 0 | 0% |

**Supplementary Table S2.** Summary the number of tetrads evaluated and their genotypic analysis

| **Diploid strain** | **No. of tetrads** | **Genotype Analysis** | | |
| --- | --- | --- | --- | --- |
|  |  | **Parental ditype** | **Tetratype** | **Non-Parental ditype** |
| W303(*fol2Δ::HIS3/FOL2 opt1Δ::LEU2/OPT1)* | 32 | 6 | 20 | 6 |

**Supplementary Table S3.** Peak appearances in positive polarity mode LC-MS/MS (Q1/Q3 parameters)

| Metabolites | Q1(Parent) | Q3(Product) | CE |
| --- | --- | --- | --- |
| Folinic acid (FN) | 512.50 | 440.60 | 28 |
| Folic acid (FA) | 442.40 | 296.30 | 28 |
| 5-methy-tetrahydrofolate (MTHF) | 498.100 | 350.10 | 28 |

**Supplementary Table S4.** List of the representative members of the OPT family used to construct the phylogenetic tree

| **Protein** | **Accession Number** | **Organism Name** | **Protein** | **Accession Number** | **Organism Name** |
| --- | --- | --- | --- | --- | --- |
| **Yeast and Fungi** | | | **Plants** | | |
| AflOpt1 | XP_071363449.1 | *Aspergillus flavus* | AtOpt1 | [NP_200404.1](https://www.ncbi.nlm.nih.gov/protein/NP_200404.1) | *Arabidopsis thaliana* |
| AfOptA | XP_752732.1 | *Aspergillus fumigatus* | AtOpt2 | [NP_172464.1](https://www.ncbi.nlm.nih.gov/protein/NP_172464.1) | *Arabidopsis thaliana* |
| AfOptB | XP_755859.1 | *Aspergillus fumigatus* | AtOpt3 | [NP_567493.5](https://www.ncbi.nlm.nih.gov/protein/NP_567493.5) | *Arabidopsis thaliana* |
| AfOptC | XP_754401.1 | *Aspergillus fumigatus* | AtOpt4 | [NP_201246.1](https://www.ncbi.nlm.nih.gov/protein/NP_201246.1) | *Arabidopsis thaliana* |
| AfOptD | XP_753269.1 | *Aspergillus fumigatus* | AtOpt5 | NP_194389.1 | *Arabidopsis thaliana* |
| AfOptE | XP_750894.1 | *Aspergillus fumigatus* | AtOpt6 | [NP_194503.1](https://www.ncbi.nlm.nih.gov/protein/NP_194503.1) | *Arabidopsis thaliana* |
| AfOptF | XP_750911.1 | *Aspergillus fumigatus* | AtOpt7 | [NP_192815.1](https://www.ncbi.nlm.nih.gov/protein/NP_192815.1) | *Arabidopsis thaliana* |
| AfOptG | XP_747762.1 | *Aspergillus fumigatus* | AtOpt8 | [NP_200164.1](https://www.ncbi.nlm.nih.gov/protein/NP_200164.1) | *Arabidopsis thaliana* |
| AfOptH | XP_746839.1 | *Aspergillus fumigatus* | AtOpt9 | [NP_200163.1](https://www.ncbi.nlm.nih.gov/protein/NP_200163.1) | *Arabidopsis thaliana* |
| AnOpt1 | XP_001397394.1 | *Aspergillus niger* | AtYsl1 | AAS00691.1 | *Arabidopsis thaliana* |
| CaOpt1 | XP_718267 | *Candida albicans* | AtYsl2 | NP_197826.2 | *Arabidopsis thaliana* |
| CaOpt2 | XP_713434 | *Candida albicans* | AtYsl3 | NP_001190532.1 | *Arabidopsis thaliana* |
| CaOpt3 | XP_713437 | *Candida albicans* | AtYsl7 | NP_176750.1 | *Arabidopsis thaliana* |
| CaOpt4 | XP_710713 | *Candida albicans* | BpeYsl | NP_879902.1 | *Bordetella pertussis* |
| CaOpt5 | XP_717657 | *Candida albicans* | MtuYsl | NP_216911.1 | *Mycobacterium tuberculosis* |
| CaOpt6 | XP_722726 | *Candida albicans* | MtYsl | WP_057365923.1 | *Mycobacterium tuberculosis* |
| CaOpt7 | XP_713354 | *Candida albicans* | MxYsl | WP_011551053.1 | *Myxococcus* |
| CaOpt8 | [XP_718105](https://sky-blast.com/accession/XP_718105) | *Candida albicans* | NmeYsl | YP_001598245.1 | *Neisseria meningitidis* |
| ClOpt1 | XP_002615150.1 | *Candida lusitaniae* | NtYsl1 | BAF48331.1 | *Nicotiana tabacum* |
| CnOpt1 | XP_772672.1 | *Cryptococcus neoformans* | OsOpt4 | XP_015638703.1 | *Oryza sativa* |
| DdYsl | XP_635289.1 | *Dictyostelium discoideum* | OsOpt5 | XP_015650551.1 | *Oryza sativa* |
| KlOpt1 | XP_453962.1 | *Kluyveromyces lactis* | OsYsl1 | BAD26556.1 | *Oryza sativa* |
| KlYsl | XP_455569.1 | *Kluyveromyces lactis* | TaOpt4 | XP_044444489.1 | *Triticum aestivum* |
| NcYsl | CAB99186.1 | *Neurospora crassa* | TaOpt7 | XP_044419492.1 | *Triticum aestivum* |
| PcOpt1 | XP_002567631.1 | *Pichia chrysogenum* | Vvi | CAN81605.1 | *Vitis vinifera* |
| PgOpt1 | XP_001486861.1 | *Pichia guilliermondii* | ZmOpt4 | NP_001152089.2 | *Zea mays* |
| Pgt1 | NP_594987.1 | *Schizosaccharomyces pombe* | ZmYsl | NP_001104952.1 | *Zea mays* |
| PpOpt1 | XP_002493413.1 | *Pichia pastoris* |  |  |  |
| ScOpt1 | NP_012323.1 | *Saccharomyces cerevisiae* | **Bacteria** | | |
| ScOpt2 | NP_015520.1 | *Saccharomyces cerevisiae* | BpeYsl | NP_879902.1 | *Bordetella pertussis* |
| ScYsl | NP_011401.1 | *Saccharomyces cerevisiae* | MtuYsl | NP_216911.1 | *Mycobacterium tuberculosis* |
| SjOpt1 | XP_002172910.1 | *Schizosaccharomyces japonicus* | MtYsl | WP_057365923.1 | *Mycobacterium tuberculosis* |
| Ssc | XP_001596456.1 | *Sclerotinia sclerotiorum* | MxYsl | WP_011551053.1 | *Myxococcus* |
| YliYsl | XP_503395.1 | *Yarrowia lipolytica* | NmeYsl | YP_001598245.1 | *Neisseria meningitidis* |

**Supplementary Table S5.** List of bacterial and yeast strains used

| **Strain** | **Genotype** | **Source** |
| --- | --- | --- |
| **Bacterial strain** | | |
| **ABE 460 DH5α** | F^+^ gyrA(Na1) recA1 endA1 thi-1 hsdR17(rm mk) GlnV44 deoRΔ (lacZYA-argF)U169 | Novagen |
| **Yeast strains** | | |
| **ABC 733** | BY4741; S288C isogenic yeast strain: MATa; his3Δ1; leu2Δ0; met15Δ0; ura3Δ0 | Euroscarf |
| **ABC734** | BY4742; S288C isogenic yeast strain: MATα; his3Δ1; leu2Δ0; lys2Δ0; ura3Δ0 | Euroscarf |
| **ABC6478** | BY4741; MATa; his3Δ1; leu2Δ0; met15Δ0; ura3Δ0; fol2Δ::HIS3Δ | This study |
| **ABC6480** | BY4742; MATα; his3Δ1; leu2Δ0; lys2Δ0; ura3Δ0; fol2Δ::HIS3Δ | This study |
| **ABC6513** | W303; MATa {leu2-3,112 trp1-1 can1-100 ura3-1 ade2-1 his3-11,15} [phi+] | Euroscarf |
| **ABC6514** | W303; MATα {leu2-3,112 trp1-1 can1-100 ura3-1 ade2-1 his3-11,15} [phi+] | Euroscarf |
| **ABC6515** | W303; MATa; {leu2-3,112 trp1-1 can1-100 ura3-1 ade2-1 his3-11,15} [phi+]; opt1Δ::LEU2Δ | This study |
| **ABC6516** | W303; MATα; {leu2-3,112 trp1-1 can1-100 ura3-1 ade2-1 his3-11,15} [phi+]; fol2Δ::HIS3Δ | This study |
| **ABC6110** | BY4741; MATa; his3Δ1; leu2Δ0; met15Δ0; ura3Δ0; YHL008c::KANMX4 | Euroscarf |
| **ABC6111** | BY4741; MATa; his3Δ1; leu2Δ0; met15Δ0; ura3Δ0; YJL193w::KANMX4 | Euroscarf |
| **ABC6106** | BY4741; MATa; his3Δ1; leu2Δ0; met15Δ0; ura3Δ0; YER039c::KANMX4 | Euroscarf |
| **ABC6107** | BY4741; MATa; his3Δ1; leu2Δ0; met15Δ0; ura3Δ0; YER060w::KANMX4 | Euroscarf |
| **ABC6108** | BY4741; MATa; his3Δ1; leu2Δ0; met15Δ0; ura3Δ0; YER060w-a::KANMX4 | Euroscarf |
| **ABC6109** | BY4741; MATa; his3Δ1; leu2Δ0; met15Δ0; ura3Δ0; YMR253c::KANMX4 | Euroscarf |
| **ABC6112** | BY4741; MATa; his3Δ1; leu2Δ0; met15Δ0; ura3Δ0; YJL163c::KANMX4 | Euroscarf |
| **ABC6113** | BY4741; MATa; his3Δ1; leu2Δ0; met15Δ0; ura3Δ0; YOR306c::KANMX4 | Euroscarf |
| **ABC6114** | BY4741; MATa; his3Δ1; leu2Δ0; met15Δ0; ura3Δ0; YOL075c:: KANMX4 | Euroscarf |
| **ABC6115** | BY4741; MATa; his3Δ1; leu2Δ0; met15Δ0; ura3Δ0; YOR071c::KANMX4 | Euroscarf |
| **ABC6117** | BY4741; MATa; his3Δ1; leu2Δ0; met15Δ0; ura3Δ0; YBL081w::KANMX4 | Euroscarf |
| **ABC6119** | BY4741; MATa; his3Δ1; leu2Δ0; met15Δ0; ura3Δ0; YBR132c::KANMX4 | Euroscarf |
| **ABC6120** | BY4741; MATa; his3Δ1; leu2Δ0; met15Δ0; ura3Δ0; YBR220c::KANMX4 | Euroscarf |
| **ABC6121** | BY4741; MATa; his3Δ1; leu2Δ0; met15Δ0; ura3Δ0; YCL049c::KANMX4 | Euroscarf |
| **ABC6122** | BY4741; MATa; his3Δ1; leu2Δ0; met15Δ0; ura3Δ0; YDL199c::KANMX4 | Euroscarf |
| **ABC6123** | BY4741; MATa; his3Δ1; leu2Δ0; met15Δ0; ura3Δ0; YGL114w::KANMX4 | Euroscarf |
| **ABC6124** | BY4741; MATa; his3Δ1; leu2Δ0; met15Δ0; ura3Δ0; YGL140c::KANMX4 | Euroscarf |
| **ABC6125** | BY4741; MATa; his3Δ1; leu2Δ0; met15Δ0; ura3Δ0; YKL217w::KANMX4 | Euroscarf |
| **ABC6126** | BY4741; MATa; his3Δ1; leu2Δ0; met15Δ0; ura3Δ0; YCR101c::KANMX4 | Euroscarf |
| **ABC6128** | BY4741; MATa; his3Δ1; leu2Δ0; met15Δ0; ura3Δ0; YNR062c::KANMX4 | Euroscarf |
| **ABC6129** | BY4742; MATα; his3Δ1; leu2Δ0; lys2Δ0; ura3Δ0; YML038c::KANMX4 | Euroscarf |
| **ABC6131** | BY4741; MATa; his3Δ1; leu2Δ0; met15Δ0; ura3Δ0; YGL186c::KANMX4 | Euroscarf |
| **ABC1611** | BY4741; MATa; his3Δ1; leu2Δ0; met15Δ0; ura3Δ0; YDR406w::KANMX4 | Euroscarf |
| **ABC1594** | BY4741; MATa; his3Δ1; leu2Δ0; met15Δ0; ura3Δ0; YOR328W::KANMX4 | Euroscarf |
| **ABC4757** | BY4741; MATa; his3Δ1; leu2Δ0; met15Δ0; ura3Δ0; YLR152C::KANMX4 | Euroscarf |
| **ABC2060** | BY4741; MATa; his3Δ1; leu2Δ0; met15Δ0; ura3Δ0; YFL055w::KANMX4 | Euroscarf |
| **ABC1530** | BY4741; MATa; his3Δ1; leu2Δ0; met15Δ0; ura3Δ0; YLL061w::KANMX4 | Euroscarf |
| **ABC1542** | BY4741; MATa; his3Δ1; leu2Δ0; met15Δ0; ura3Δ0; YPL274w::KANMX4 | Euroscarf |
| **ABC4467** | BY4741; MATa; his3Δ1; leu2Δ0; met15Δ0; ura3Δ0; YNL 125c::KANMX4 | Euroscarf |
| **ABC4444** | BY4741; MATa; his3Δ1; leu2Δ0; met15Δ0; ura3Δ0; YOL119C::KANMX4 | Euroscarf |
| **ABC2067** | BY4741; MATa; his3Δ1; leu2Δ0; met15Δ0; ura3Δ0; YAL067c::KANMX4 | Euroscarf |
| **ABC2061** | BY4742; MATα; his3Δ1; leu2Δ0; lys2Δ0; ura3Δ0; YGR260w::KANMX4 | Euroscarf |
| **ABC2073** | BY4741; MATa; his3Δ1; leu2Δ0; met15Δ0; ura3Δ0;YLR004c::KANMX4 | Euroscarf |
| **ABC1651** | BY4741; MATa; his3Δ1; leu2Δ0; met15Δ0; ura3Δ0;YOL162w::KANMX4 | Euroscarf |
| **ABC1631** | BY4741; MATa; his3Δ1; leu2Δ0; met15Δ0; ura3Δ0; YOL163w::KANMX4 | Euroscarf |
| **ABC2354** | BY4741; MATa; his3Δ1; leu2Δ0; met15Δ0; ura3Δ0; YIR028W:: KANMX4 | Euroscarf |
| **ABC6389** | BY4741; MATa; his3Δ1; leu2Δ0; met15Δ0; ura3Δ0; YCR011c::kanMX4 | Euroscarf |
| **ABC6390** | BY4741; MATa; his3Δ1; leu2Δ0; met15Δ0; ura3Δ0; YKR104w::kanMX4 | Euroscarf |
| **ABC6391** | BY4741; MATa; his3Δ1; leu2Δ0; met15Δ0; ura3Δ0; YKR103w::kanMX4 | Euroscarf |
| **ABC6392** | BY4741; MATa; his3Δ1; leu2Δ0; met15Δ0; ura3Δ0; YPR201w::kanMX4 | Euroscarf |
| **ABC6393** | BY4741; MATa; his3Δ1; leu2Δ0; met15Δ0; ura3Δ0; YDL206w::kanMX4 | Euroscarf |
| **ABC6394** | BY4741; MATa; his3Δ1; leu2Δ0; met15Δ0; ura3Δ0; YDR387c::kanMX4 | Euroscarf |
| **ABC6395** | BY4741; MATa; his3Δ1; leu2Δ0; met15Δ0; ura3Δ0; YFL040w::kanMX4 | Euroscarf |
| **ABC6396** | BY4741; MATa; his3Δ1; leu2Δ0; met15Δ0; ura3Δ0; YKR105c::kanMX4 | Euroscarf |
| **ABC6397** | BY4741; MATa; his3Δ1; leu2Δ0; met15Δ0; ura3Δ0; YHR048w::kanMX4 | Euroscarf |
| **ABC6398** | BY4741; MATa; his3Δ1; leu2Δ0; met15Δ0; ura3Δ0; YNR055c::kanMX4 | Euroscarf |
| **ABC6399** | BY4741; MATa; his3Δ1; leu2Δ0; met15Δ0; ura3Δ0; YCR028c::kanMX4 | Euroscarf |
| **ABC6401** | BY4741; MATa; his3Δ1; leu2Δ0; met15Δ0; ura3Δ0; YBR219c::kanMX4 | Euroscarf |
| **ABC6402** | BY4741; MATa; his3Δ1; leu2Δ0; met15Δ0; ura3Δ0; YDL054c::kanMX4 | Euroscarf |
| **ABC6403** | BY4741; MATa; his3Δ1; leu2Δ0; met15Δ0; ura3Δ0; YOL137w::kanMX4 | Euroscarf |
| **ABC6404** | BY4741; MATa; his3Δ1; leu2Δ0; met15Δ0; ura3Δ0; YLR237w::kanMX4 | Euroscarf |
| **ABC6405** | BY4741; MATa; his3Δ1; leu2Δ0; met15Δ0; ura3Δ0; YOR192c::kanMX4 | Euroscarf |
| **ABC6406** | BY4741; MATa; his3Δ1; leu2Δ0; met15Δ0; ura3Δ0; YLR034c::kanMX4 | Euroscarf |
| **ABC6407** | BY4741; MATa; his3Δ1; leu2Δ0; met15Δ0; ura3Δ0; YJL107c::kanMX4 | Euroscarf |
| **ABC6408** | BY4741; MATa; his3Δ1; leu2Δ0; met15Δ0; ura3Δ0; YMR319c::kanMX4 | Euroscarf |
| **ABC6409** | BY4741; MATa; his3Δ1; leu2Δ0; met15Δ0; ura3Δ0; YMR155w::kanMX4 | Euroscarf |
| **ABC6410** | BY4741; MATa; his3Δ1; leu2Δ0; met15Δ0; ura3Δ0; YJR124c::kanMX4 | Euroscarf |
| **ABC6411** | BY4741; MATa; his3Δ1; leu2Δ0; met15Δ0; ura3Δ0; YBR008c::kanMX4 | Euroscarf |
| **ABC6412** | BY4741; MATa; his3Δ1; leu2Δ0; met15Δ0; ura3Δ0; YIL134w::kanMX4 | Euroscarf |
| **ABC6413** | BY4741; MATa; his3Δ1; leu2Δ0; met15Δ0; ura3Δ0; YML047c::kanMX4 | Euroscarf |
| **ABC6414** | BY4741; MATa; his3Δ1; leu2Δ0; met15Δ0; ura3Δ0; YFL011w::kanMX4 | Euroscarf |
| **ABC6415** | BY4741; MATa; his3Δ1; leu2Δ0; met15Δ0; ura3Δ0; YBL049w::kanMX4 | Euroscarf |
| **ABC6416** | BY4741; MATa; his3Δ1; leu2Δ0; met15Δ0; ura3Δ0; YKL221w::kanMX4 | Euroscarf |
| **ABC6417** | BY4741; MATa; his3Δ1; leu2Δ0; met15Δ0; ura3Δ0; YHR096c::kanMX4 | Euroscarf |
| **ABC6418** | BY4741; MATa; his3Δ1; leu2Δ0; met15Δ0; ura3Δ0; YGR138c::kanMX4 | Euroscarf |
| **ABC6419** | BY4741; MATa; his3Δ1; leu2Δ0; met15Δ0; ura3Δ0; YPR192w::kanMX4 | Euroscarf |
| **ABC6420** | BY4741; MATa; his3Δ1; leu2Δ0; met15Δ0; ura3Δ0; YFL054c::kanMX4 | Euroscarf |
| **ABC6421** | BY4741; MATa; his3Δ1; leu2Δ0; met15Δ0; ura3Δ0; YOL084w::kanMX4 | Euroscarf |
| **ABC6422** | BY4741; MATa; his3Δ1; leu2Δ0; met15Δ0; ura3Δ0; YLL005c::kanMX4 | Euroscarf |
| **ABC6423** | BY4741; MATa; his3Δ1; leu2Δ0; met15Δ0; ura3Δ0; YDR536w::kanMX4 | Euroscarf |
| **ABC6424** | BY4741; MATa; his3Δ1; leu2Δ0; met15Δ0; ura3Δ0; YKR106w::kanMX4 | Euroscarf |
| **ABC1092** | BY4742; MATα; his3Δ1; leu2Δ0; lys2Δ0; ura3Δ0; YGL255W::kanMX4 | Euroscarf |
| **ABC2178** | BY4741; MAT; his3Δ1; leu2Δ0; met15Δ0; ura3Δ0; YPL265W::kanMX4 | Euroscarf |
| **ABC4442** | BY4741; MATa; his3Δ1; leu2Δ0; met15Δ0; ura3Δ0; YNR070W::kanMX4 | Euroscarf |
| **ABC1733** | BY4742; MATα; his3Δ1; leu2Δ0; lys2Δ0; ura3Δ0; YPR124W::kanMX4 | Euroscarf |
| **ABC1530** | BY4741; MATa; his3Δ1; leu2Δ0; met15Δ0; ura3Δ0; YLL061W::kanMX4 | Euroscarf |
| **ABC4385** | BY4741; MATa; his3Δ1; leu2Δ0; met15Δ0; ura3Δ0; YCR010C::kanMX4 | Euroscarf |
| **ABC1844** | BY4741; MATa; his3Δ1; leu2Δ0; met15Δ0; ura3Δ0; YHL036W::kanMX4 | Euroscarf |
| **ABC2174** | BY4741; MATa; his3Δ1; leu2Δ0; met15Δ0; ura3Δ0; YOR348C::kanMX4 | Euroscarf |
| **ABC1481** | BY4741; MATa; his3Δ1; leu2Δ0; met15Δ0; ura3Δ0; YJL212C::kanMX4 | Euroscarf |
| **ABC6327** | BY4741; MATa; his3Δ1; leu2Δ0; met15Δ0; ura3Δ0; YHL008c::KANMX4; fol2Δ::HIS3Δ | This study |
| **ABC6328** | BY4741; MATa; his3Δ1; leu2Δ0; met15Δ0; ura3Δ0; YJL193w::KANMX4; fol2Δ::HIS3Δ | This study |
| **ABC6323** | BY4741; MATa; his3Δ1; leu2Δ0; met15Δ0; ura3Δ0; YER039c::KANMX4; fol2Δ::HIS3Δ | This study |
| **ABC6324** | BY4741; MATa; his3Δ1; leu2Δ0; met15Δ0; ura3Δ0; YER060w::KANMX4; fol2Δ::HIS3Δ | This study |
| **ABC6325** | BY4741; MATa; his3Δ1; leu2Δ0; met15Δ0; ura3Δ0; YER060w-a::KANMX4; fol2Δ::HIS3Δ | This study |
| **ABC6326** | BY4741; MATa; his3Δ1; leu2Δ0; met15Δ0; ura3Δ0; YMR253c::KANMX4; fol2Δ::HIS3Δ | This study |
| **ABC6329** | BY4741; MATa; his3Δ1; leu2Δ0; met15Δ0; ura3Δ0; YJL163c::KANMX4; fol2Δ::HIS3Δ | This study |
| **ABC6330** | BY4741; MATa; his3Δ1; leu2Δ0; met15Δ0; ura3Δ0; YOR306c::KANMX4; fol2Δ::HIS3Δ | This study |
| **ABC6331** | BY4741; MATa; his3Δ1; leu2Δ0; met15Δ0; ura3Δ0; YOL075c:: KANMX4; fol2Δ::HIS3Δ | This study |
| **ABC6332** | BY4741; MATa; his3Δ1; leu2Δ0; met15Δ0; ura3Δ0; YOR071c::KANMX4; fol2Δ::HIS3Δ | This study |
| **ABC6333** | BY4741; MATa; his3Δ1; leu2Δ0; met15Δ0; ura3Δ0; YBL081w::KANMX4; fol2Δ::HIS3Δ | This study |
| **ABC6334** | BY4741; MATa; his3Δ1; leu2Δ0; met15Δ0; ura3Δ0; YBR132c::KANMX4; fol2Δ::HIS3Δ | This study |
| **ABC6335** | BY4741; MATa; his3Δ1; leu2Δ0; met15Δ0; ura3Δ0; YBR220c::KANMX4; fol2Δ::HIS3Δ | This study |
| **ABC6336** | BY4741; MATa; his3Δ1; leu2Δ0; met15Δ0; ura3Δ0; YCL049c::KANMX4; fol2Δ::HIS3Δ | This study |
| **ABC6337** | BY4741; MATa; his3Δ1; leu2Δ0; met15Δ0; ura3Δ0; YDL199c::KANMX4; fol2Δ::HIS3Δ | This study |
| **ABC6338** | BY4741; MATa; his3Δ1; leu2Δ0; met15Δ0; ura3Δ0; YGL114w::KANMX4; fol2Δ::HIS3Δ | This study |
| **ABC6339** | BY4741; MATa; his3Δ1; leu2Δ0; met15Δ0; ura3Δ0; YGL140c::KANMX4; fol2Δ::HIS3Δ | This study |
| **ABC6340** | BY4741; MATa; his3Δ1; leu2Δ0; met15Δ0; ura3Δ0; YKL217w::KANMX4; fol2Δ::HIS3Δ | This study |
| **ABC6341** | BY4741; MATa; his3Δ1; leu2Δ0; met15Δ0; ura3Δ0; YCR101c::KANMX4; fol2Δ::HIS3Δ | This study |
| **ABC6342** | BY4741; MATa; his3Δ1; leu2Δ0; met15Δ0; ura3Δ0; YNR062c::KANMX4; fol2Δ::HIS3Δ | This study |
| **ABC6343** | BY4742; MATα; his3Δ1; leu2Δ0; lys2Δ0; ura3Δ0; YML038c::KANMX4; fol2Δ::HIS3Δ | This study |
| **ABC6344** | BY4741; MATa; his3Δ1; leu2Δ0; met15Δ0; ura3Δ0; YGL186c::KANMX4; fol2Δ::HIS3Δ | This study |
| **ABC6345** | BY4741; MATa; his3Δ1; leu2Δ0; met15Δ0; ura3Δ0; YDR406w::KANMX4; fol2Δ::HIS3Δ | This study |
| **ABC6346** | BY4741; MATa; his3Δ1; leu2Δ0; met15Δ0; ura3Δ0; YOR328W::KANMX4; fol2Δ::HIS3Δ | This study |
| **ABC6347** | BY4741; MATa; his3Δ1; leu2Δ0; met15Δ0; ura3Δ0; YLR152C::KANMX4; fol2Δ::HIS3Δ | This study |
| **ABC6348** | BY4741; MATa; his3Δ1; leu2Δ0; met15Δ0; ura3Δ0; YFL055w::KANMX4; fol2Δ::HIS3Δ | This study |
| **ABC6349** | BY4741; MATa; his3Δ1; leu2Δ0; met15Δ0; ura3Δ0; YLL061w::KANMX4; fol2Δ::HIS3Δ | This study |
| **ABC6350** | BY4741; MATa; his3Δ1; leu2Δ0; met15Δ0; ura3Δ0; YPL274w::KANMX4; fol2Δ::HIS3Δ | This study |
| **ABC6351** | BY4741; MATa; his3Δ1; leu2Δ0; met15Δ0; ura3Δ0; YNL 125c::KANMX4; fol2Δ::HIS3Δ | This study |
| **ABC6352** | BY4741; MATa; his3Δ1; leu2Δ0; met15Δ0; ura3Δ0; YOL119C::KANMX4; fol2Δ::HIS3Δ | This study |
| **ABC6353** | BY4741; MATa; his3Δ1; leu2Δ0; met15Δ0; ura3Δ0; YAL067c::KANMX4; fol2Δ::HIS3Δ | This study |
| **ABC6354** | BY4742; MATα; his3Δ1; leu2Δ0; lys2Δ0; ura3Δ0; YGR260w::KANMX4; fol2Δ::HIS3Δ | This study |
| **ABC6355** | BY4741; MATa; his3Δ1; leu2Δ0; met15Δ0; ura3Δ0;YLR004c::KANMX4; fol2Δ::HIS3Δ | This study |
| **ABC6356** | BY4741; MATa; his3Δ1; leu2Δ0; met15Δ0; ura3Δ0;YOL162w::KANMX4; fol2Δ::HIS3Δ | This study |
| **ABC6357** | BY4741; MATa; his3Δ1; leu2Δ0; met15Δ0; ura3Δ0; YOL163w::KANMX4; fol2Δ::HIS3Δ | This study |
| **ABC6358** | BY4741; MATa; his3Δ1; leu2Δ0; met15Δ0; ura3Δ0; YIR028W:: KANMX4; fol2Δ::HIS3Δ | This study |
| **ABC6468** | BY4741; MATa; his3Δ1; leu2Δ0; met15Δ0; ura3Δ0; YCR011c::kanMX4; fol2Δ::HIS3Δ | This study |
| **ABC6469** | BY4741; MATa; his3Δ1; leu2Δ0; met15Δ0; ura3Δ0; YKR104w::kanMX4; fol2Δ::HIS3Δ | This study |
| **ABC6470** | BY4741; MATa; his3Δ1; leu2Δ0; met15Δ0; ura3Δ0; YKR103w::kanMX4; fol2Δ::HIS3Δ | This study |
| **ABC6471** | BY4741; MATa; his3Δ1; leu2Δ0; met15Δ0; ura3Δ0; YPR201w::kanMX4; fol2Δ::HIS3Δ | This study |
| **ABC6435** | BY4741; MATa; his3Δ1; leu2Δ0; met15Δ0; ura3Δ0; YDL206w::kanMX4; fol2Δ::HIS3Δ | This study |
| **ABC6436** | BY4741; MATa; his3Δ1; leu2Δ0; met15Δ0; ura3Δ0; YDR387c::kanMX4; fol2Δ::HIS3Δ | This study |
| **ABC6437** | BY4741; MATa; his3Δ1; leu2Δ0; met15Δ0; ura3Δ0; YFL040w::kanMX4; fol2Δ::HIS3Δ | This study |
| **ABC6438** | BY4741; MATa; his3Δ1; leu2Δ0; met15Δ0; ura3Δ0; YKR105c::kanMX4; fol2Δ::HIS3Δ | This study |
| **ABC6439** | BY4741; MATa; his3Δ1; leu2Δ0; met15Δ0; ura3Δ0; YHR048w::kanMX4; fol2Δ::HIS3Δ | This study |
| **ABC6440** | BY4741; MATa; his3Δ1; leu2Δ0; met15Δ0; ura3Δ0; YNR055c::kanMX4; fol2Δ::HIS3Δ | This study |
| **ABC6441** | BY4741; MATa; his3Δ1; leu2Δ0; met15Δ0; ura3Δ0; YCR028c::kanMX4; fol2Δ::HIS3Δ | This study |
| **ABC6443** | BY4741; MATa; his3Δ1; leu2Δ0; met15Δ0; ura3Δ0; YBR219c::kanMX4; fol2Δ::HIS3Δ | This study |
| **ABC6444** | BY4741; MATa; his3Δ1; leu2Δ0; met15Δ0; ura3Δ0; YDL054c::kanMX4; fol2Δ::HIS3Δ | This study |
| **ABC6445** | BY4741; MATa; his3Δ1; leu2Δ0; met15Δ0; ura3Δ0; YOL137w::kanMX4; fol2Δ::HIS3Δ | This study |
| **ABC6446** | BY4741; MATa; his3Δ1; leu2Δ0; met15Δ0; ura3Δ0; YLR237w::kanMX4; fol2Δ::HIS3Δ | This study |
| **ABC6447** | BY4741; MATa; his3Δ1; leu2Δ0; met15Δ0; ura3Δ0; YOR192c::kanMX4; fol2Δ::HIS3Δ | This study |
| **ABC6448** | BY4741; MATa; his3Δ1; leu2Δ0; met15Δ0; ura3Δ0; YLR034c::kanMX4; fol2Δ::HIS3Δ | This study |
| **ABC6449** | BY4741; MATa; his3Δ1; leu2Δ0; met15Δ0; ura3Δ0; YJL107c::kanMX4; fol2Δ::HIS3Δ | This study |
| **ABC6450** | BY4741; MATa; his3Δ1; leu2Δ0; met15Δ0; ura3Δ0; YMR319c::kanMX4; fol2Δ::HIS3Δ | This study |
| **ABC6451** | BY4741; MATa; his3Δ1; leu2Δ0; met15Δ0; ura3Δ0; YMR155w::kanMX4; fol2Δ::HIS3Δ | This study |
| **ABC6452** | BY4741; MATa; his3Δ1; leu2Δ0; met15Δ0; ura3Δ0; YJR124c::kanMX4; fol2Δ::HIS3Δ | This study |
| **ABC6453** | BY4741; MATa; his3Δ1; leu2Δ0; met15Δ0; ura3Δ0; YBR008c::kanMX4; fol2Δ::HIS3Δ | This study |
| **ABC6454** | BY4741; MATa; his3Δ1; leu2Δ0; met15Δ0; ura3Δ0; YIL134w::kanMX4; fol2Δ::HIS3Δ | This study |
| **ABC6455** | BY4741; MATa; his3Δ1; leu2Δ0; met15Δ0; ura3Δ0; YML047c::kanMX4; fol2Δ::HIS3Δ | This study |
| **ABC6456** | BY4741; MATa; his3Δ1; leu2Δ0; met15Δ0; ura3Δ0; YFL011w::kanMX4; fol2Δ::HIS3Δ | This study |
| **ABC6457** | BY4741; MATa; his3Δ1; leu2Δ0; met15Δ0; ura3Δ0; YBL049w::kanMX4; fol2Δ::HIS3Δ | This study |
| **ABC6458** | BY4741; MATa; his3Δ1; leu2Δ0; met15Δ0; ura3Δ0; YKL221w::kanMX4; fol2Δ::HIS3Δ | This study |
| **ABC6459** | BY4741; MATa; his3Δ1; leu2Δ0; met15Δ0; ura3Δ0; YHR096c::kanMX4; fol2Δ::HIS3Δ | This study |
| **ABC6460** | BY4741; MATa; his3Δ1; leu2Δ0; met15Δ0; ura3Δ0; YGR138c::kanMX4; fol2Δ::HIS3Δ | This study |
| **ABC6461** | BY4741; MATa; his3Δ1; leu2Δ0; met15Δ0; ura3Δ0; YPR192w::kanMX4; fol2Δ::HIS3Δ | This study |
| **ABC6462** | BY4741; MATa; his3Δ1; leu2Δ0; met15Δ0; ura3Δ0; YFL054c::kanMX4; fol2Δ::HIS3Δ | This study |
| **ABC6463** | BY4741; MATa; his3Δ1; leu2Δ0; met15Δ0; ura3Δ0; YOL084w::kanMX4; fol2Δ::HIS3Δ | This study |
| **ABC6464** | BY4741; MATa; his3Δ1; leu2Δ0; met15Δ0; ura3Δ0; YLL005c::kanMX4; fol2Δ::HIS3Δ | This study |
| **ABC6465** | BY4741; MATa; his3Δ1; leu2Δ0; met15Δ0; ura3Δ0; YDR536w::kanMX4; fol2Δ::HIS3Δ | This study |
| **ABC6466** | BY4741; MATa; his3Δ1; leu2Δ0; met15Δ0; ura3Δ0; YKR106w::kanMX4; fol2Δ::HIS3Δ | This study |
| **ABC6359** | BY4742; MATα; his3Δ1; leu2Δ0; lys2Δ0; ura3Δ0; YGL255W::kanMX4; fol2Δ::HIS3Δ | This study |
| **ABC6360** | BY4741; MAT; his3Δ1; leu2Δ0; met15Δ0; ura3Δ0; YPL265W::kanMX4; fol2Δ::HIS3Δ | This study |
| **ABC6361** | BY4741; MATa; his3Δ1; leu2Δ0; met15Δ0; ura3Δ0; YNR070W::kanMX4; fol2Δ::HIS3Δ | This study |
| **ABC6362** | BY4742; MATα; his3Δ1; leu2Δ0; lys2Δ0; ura3Δ0; YPR124W::kanMX4; fol2Δ::HIS3Δ | This study |
| **ABC6363** | BY4741; MATa; his3Δ1; leu2Δ0; met15Δ0; ura3Δ0; YLL061W::kanMX4; fol2Δ::HIS3Δ | This study |
| **ABC6364** | BY4741; MATa; his3Δ1; leu2Δ0; met15Δ0; ura3Δ0; YCR010C::kanMX4; fol2Δ::HIS3Δ | This study |
| **ABC6365** | BY4741; MATa; his3Δ1; leu2Δ0; met15Δ0; ura3Δ0; YHL036W::kanMX4; fol2Δ::HIS3Δ | This study |
| **ABC6366** | BY4741; MATa; his3Δ1; leu2Δ0; met15Δ0; ura3Δ0; YOR348C::kanMX4; fol2Δ::HIS3Δ | This study |

**Supplementary Table S6.**  List of plasmids used in the study

| **Plasmid name** | **Clone No** | **Description** | **Source** |
| --- | --- | --- | --- |
| **p416TEF** | ABE 443 | Centromeric shuttle vector bearing URA3 marker, TEF promoter, MCS region, and CYC terminator for yeast expression and Amprmarker for selection in E. coli | Dr. Martin Funk |
| **pDR_AtOPT3** | ABE6548 | AtOPT3 cloned in vector pDR | Dr. Cecile Gaillard |
| **pDR_AtOPT6** | ABE6550 | AtOPT6 cloned in vector pDR | Dr. Cecile Gaillard |
| **U21422** | ABE6564 | AtOPT1 cDNA cloned in pUNI51 vector | Arabidopsis Biological Resource Center (ABRC) |
| **DQ446240)** | ABE6565 | AtOPT2 cloned in the Gateway(TM) pDONR221 recombination vector | Arabidopsis Biological Resource Center (ABRC) |
| **U25639** | ABE6566 | AtOPT4 cDNA cloned in pENTR/SD-dTopo vector | Arabidopsis Biological Resource Center (ABRC) |
| **U09285** | ABE6567 | AtOPT5 cDNA cloned in pUNI51 vector | Arabidopsis Biological Resource Center (ABRC) |
| **U12998** | ABE6568 | AtOPT7 cDNA cloned in pUNI51 vector | Arabidopsis Biological Resource Center (ABRC) |
| **DQ447070** | ABE6569 | AtOPT8 cloned is in the Gateway(TM) pENTR221 recombination vector | Arabidopsis Biological Resource Center (ABRC) |
| **pSK528x** | ABE6571 | pSK528x A. fumigatus optA cDNA in pSK51 (SalI) | Dr. Sven Krappmann |
| **pSK529x** | ABE6572 | pSK529x A. fumigatus optB cDNA in pSK51 (SalI/NotI) | Dr. Sven Krappmann |
| **pSK530x** | ABE6573 | pSK530x A. fumigatus optC cDNA in pSK51 (SalI/NotI) | Dr. Sven Krappmann |
| **pSK531x** | ABE6574 | pSK531x A. fumigatus optD cDNA in pSK51 (NotI) | Dr. Sven Krappmann |
| **pSK532x** | ABE6575 | pSK532x A. fumigatus optE cDNA in pSK51 (SalI/NotI) | Dr. Sven Krappmann |
| **pSK533x** | ABE6576 | pSK533x A. fumigatus optF cDNA in pSK51 (SalI/NotI) | Dr. Sven Krappmann |
| **pSK534x** | ABE6577 | pSK534x A. fumigatus optG cDNA in pSK51 (SalI/NotI) | Dr. Sven Krappmann |
| **pSK535x** | ABE6578 | pSK535x A. fumigatus optH cDNA in pSK51 (SalI/NotI) | Dr. Sven Krappmann |
| **p416-TEF AtOPT1** | ABE6588 | AtOPT1 cDNA PCR amplified from ABE6564 and cloned in p416TEF vector using homologous recombination; BamHI and XhoI sites provided upstream and dowstream of the ORF for subcloning | This study |
| **p416-TEF AtOPT2** | ABE6589 | AtOPT2 PCR amplified from ABE6565 and cloned in p416TEF vector using homologous recombination; BamHI and XhoI sites provided upstream and dowstream of the ORF for subcloning | This study |
| **p416-TEF AtOPT3** | ABE6590 | AtOPT3 cDNA PCR amplified from ABE6548 and cloned in p416TEF vector using homologous recombination; XbaI/EcoRI and XhoI sites provided upstream and dowstream of the ORF for subcloning | This study |
| **p416-TEF AtOPT4** | ABE6591 | AtOPT4 cDNA PCR amplified from ABE6566 and cloned in p416TEF vector using homologous recombination; XbaI/EcoRI and XhoI sites provided upstream and dowstream of the ORF for subcloning | This study |
| **p416-TEF AtOPT5** | ABE6592 | AtOPT5 cDNA PCR amplified from ABE6567 and cloned in p416TEF vector using homologous recombination; XbaI/BamHI and XhoI sites provided upstream and dowstream of the ORF for subcloning | This study |
| **p416-TEF AtOPT6** | ABE6593 | AtOPT6 cDNA PCR amplified from ABE6550 and cloned in p416TEF vector using homologous recombination; XbaI/BamHI and XhoI sites provided upstream and dowstream of the ORF for subcloning | This study |
| **p416-TEF AtOPT7** | ABE6584 | AtOPT7 cDNA PCR amplified from ABE6568 and cloned in p416TEF vector using homologous recombination; XbaI/BamHI and XhoI sites provided upstream and dowstream of the ORF for subcloning | This study |
| **p416-TEF AtOPT8** | ABE6594 | AtOPT8 cDNA PCR amplified from ABE6569 and cloned in p416TEF vector using homologous recombination; XbaI/BamHI and XhoI sites provided upstream and dowstream of the ORF for subcloning | This study |
| **pRS416TEF FOL2** | ABE6246 | fol2 gene along with 321 5' overhang and 100 nt 3' overhang cloned in pRS416Tef using BamHI and XhoI | This study |
| **pRS416 FOL2::HIS3** | ABE6253 | Histidine marker cloned in pRS416FOL2::HIS3 at EcoRI site | This study |
| **p416-TEF-CaOPT7** | AB3071 | The gene encoding the putative OPT1 ortholog from C. albicans was PCR amplified using C. albicans genomic DNA and cloned in pRS416TEF vector at BamHI and XbaI stites | Lab stock |
| **p416-TEF-CaOPT1** | AB 1786 | The gene encoding the putative OPT1 ortholog from C. albicans was PCR amplified using C. albicans genomic DNA and cloned in pRS416TEF vector at EcoR1 and XhoI stites | Lab stock |
| **Mutants** |  |  |  |
| **TMD3** |  |  |  |
| **pA179G** | ABE 3850 | p416-TEF-His-HGT1-HA with A179G mutation | Lab stock |
| **pV180A** | ABE 3851 | p416-TEF-His-HGT1-HA with V180A mutation | Lab stock |
| **pI181A** | ABE 4057 | p416-TEF-His-HGT1-HA with I181A mutation | Lab stock |
| **pT182A** | ABE 3659 | p416-TEF-His-HGT1-HA with T182A mutation | Lab stock |
| **pI183A** | ABE 3852 | p416-TEF-His-HGT1-HA with I183A mutation | Lab stock |
| **pA184G** | ABE 3853 | p416-TEF-His-HGT1-HA with A184G mutation | Lab stock |
| **pV185A** | ABE 3854 | p416-TEF-His-HGT1-HA with V185A mutation | Lab stock |
| **pA186G** | ABE 3855 | p416-TEF-His-HGT1-HA with A186G mutation | Lab stock |
| **pL187A** | ABE 3856 | p416-TEF-His-HGT1-HA with L187A mutation | Lab stock |
| **pT188A** | ABE 2039 | p416-TEF-His-HGT1-HA with T188A mutation | Lab stock |
| **pS189A** | ABE 3660 | p416-TEF-His-HGT1-HA with S189A mutation | Lab stock |
| **pS190A** | ABE 2373 | p416-TEF-His-HGT1-HA with S190A mutation | Lab stock |
| **pA192G** | ABE 3857 | p416-TEF-His-HGT1-HA with A192G mutation | Lab stock |
| **pY193A** | ABE 3858 | p416-TEF-His-HGT1-HA with Y193A mutation | Lab stock |
| **pA194G** | ABE 3859 | p416-TEF-His-HGT1-HA with A194G mutation | Lab stock |
| **pM195A** | ABE 3860 | p416-TEF-His-HGT1-HA with M195A mutation | Lab stock |
| **pI197A** | ABE 3861 | p416-TEF-His-HGT1-HA with I197A mutation | Lab stock |
| **pL198A** | ABE 3862 | p416-TEF-His-HGT1-HA with L198A mutation | Lab stock |
| **pN199A** | ABE 3664 | p416-TEF-His-HGT1-HA with N199A mutation | Lab stock |
| **pA200G** | ABE 3863 | p416-TEF-His-HGT1-HA with A200G mutation | Lab stock |
| **TMD4** |  |  |  |
| **pG212A** | ABE 3192 | p416-TEF-His-HGT1-HA with G212A mutation | Lab stock |
| **pY213A** | ABE 3193 | p416-TEF-His-HGT1-HA with Y213A mutation | Lab stock |
| **pQ214A** | ABE 3082 | p416-TEF-His-HGT1-HA with Q214A mutation | Lab stock |
| **pF215A** | ABE 3194 | p416-TEF-His-HGT1-HA with F215A mutation | Lab stock |
| **pL216A** | ABE 3195 | p416-TEF-His-HGT1-HA with L216A mutation | Lab stock |
| **pV218A** | ABE 3198 | p416-TEF-His-HGT1-HA with V218A mutation | Lab stock |
| **pW219A** | ABE 3197 | p416-TEF-His-HGT1-HA with W219A mutation | Lab stock |
| **pT220A** | ABE 3191 | p416-TEF-His-HGT1-HA with T220A mutation | Lab stock |
| **pS221A** | ABE 3347 | p416-TEF-His-HGT1-HA with S221A mutation | Lab stock |
| **pQ222A** | ABE 2041 | p416-TEF-His-HGT1-HA with Q222A mutation *(Kaur et al., 2009)* | Lab stock |
| **pM223A** | ABE 3508 | p416-TEF-His-HGT1-HA with M223A mutation | Lab stock |
| **pL224A** | ABE 3448 | p416-TEF-His-HGT1-HA with L224A mutation | Lab stock |
| **pG225A** | ABE 3449 | p416-TEF-His-HGT1-HA with G225A mutation | Lab stock |
| **pY226A** | ABE 3450 | p416-TEF-His-HGT1-HA with Y226A mutation | Lab stock |
| **pG227A** | ABE 3451 | p416-TEF-His-HGT1-HA with G227A mutation | Lab stock |
| **pA228A** | ABE 3452 | p416-TEF-His-HGT1-HA with A228A mutation | Lab stock |
| **pA229C** | ABE 3453 | p416-TEF-His-HGT1-HA with A229C mutation | Lab stock |
| **pG230A** | ABE 3454 | p416-TEF-His-HGT1-HA with G230A mutation | Lab stock |
| **pL231A** | ABE 3509 | p416-TEF-His-HGT1-HA with L231A mutation | Lab stock |
| **pT232A** | ABE 3510 | p416-TEF-His-HGT1-HA with T232A mutation | Lab stock |
| **TMD5** |  |  |  |
| **pF277A** | ABE 3711 | p416-TEF-His-HGT1-HA with F277A mutation | Lab stock |
| **pF278A** | ABE 3712 | p416-TEF-His-HGT1-HA with F278A mutation | Lab stock |
| **pL279A** | ABE 3713 | p416-TEF-His-HGT1-HA with L279A mutation | Lab stock |
| **pI280A** | ABE 3714 | p416-TEF-His-HGT1-HA with I280A mutation | Lab stock |
| **pV281A** | ABE 3715 | p416-TEF-His-HGT1-HA with V281A mutation | Lab stock |
| **pL282A** | ABE 3716 | p416-TEF-His-HGT1-HA with L282A mutation | Lab stock |
| **pI283A** | ABE 3717 | p416-TEF-His-HGT1-HA with I283A mutation | Lab stock |
| **pG284A** | ABE 3718 | p416-TEF-His-HGT1-HA with G284A mutation | Lab stock |
| **pS285A** | ABE 3075 | p416-TEF-His-HGT1-HA with S285A mutation | Lab stock |
| **pF286A** | ABE 3955 | p416-TEF-His-HGT1-HA with F286A mutation | Lab stock |
| **pI287A** | ABE 3956 | p416-TEF-His-HGT1-HA with I287A mutation | Lab stock |
| **pW288A** | ABE 3719 | p416-TEF-His-HGT1-HA with W288A mutation | Lab stock |
| **pY289A** | ABE 3720 | p416-TEF-His-HGT1-HA with Y289A mutation | Lab stock |
| **pW290A** | ABE 3721 | p416-TEF-His-HGT1-HA with W290A mutation | Lab stock |
| **pV291A** | ABE 3722 | p416-TEF-His-HGT1-HA with V291A mutation | Lab stock |
| **pP292A** | ABE 2852 | p416-TEF-His-HGT1-HA with P292A mutation | Lab stock |
| **pG293A** | ABE 3723 | p416-TEF-His-HGT1-HA with G293A mutation | Lab stock |
| **pF294A** | ABE 3724 | p416-TEF-His-HGT1-HA with F294A mutation | Lab stock |
| **pL295A** | ABE 3725 | p416-TEF-His-HGT1-HA with L295A mutation | Lab stock |
| **pF296A** | ABE 3726 | p416-TEF-His-HGT1-HA with F296A mutation | Lab stock |
| **TMD6** |  |  |  |
| **pV354A** | ABE 4054 | p416-TEF-His-HGT1-HA with V354A mutation | Lab stock |
| **pA356G** | ABE 4056 | p416-TEF-His-HGT1-HA with A356G mutation | Lab stock |
| **pT358A** | ABE 3661 | p416-TEF-His-HGT1-HA with T358A mutation | Lab stock |
| **pY359A** | ABE 3880 | p416-TEF-His-HGT1-HA with Y359A mutation | Lab stock |
| **pA360G** | ABE 3881 | p416-TEF-His-HGT1-HA with A360G mutation | Lab stock |
| **pV362A** | ABE 3882 | p416-TEF-His-HGT1-HA with V362A mutation | Lab stock |
| **pL363A** | ABE 3883 | p416-TEF-His-HGT1-HA with L363A mutation | Lab stock |
| **pI364A** | ABE 3884 | p416-TEF-His-HGT1-HA with I364A mutation | Lab stock |
| **pF365A** | ABE 4058 | p416-TEF-His-HGT1-HA with F365A mutation | Lab stock |
| **pF366A** | ABE 3885 | p416-TEF-His-HGT1-HA with F366A mutation | Lab stock |
| **pV367A** | ABE 3886 | p416-TEF-His-HGT1-HA with V367A mutation | Lab stock |
| **pI368A** | ABE 4059 | p416-TEF-His-HGT1-HA with I368A mutation | Lab stock |
| **pL370A** | ABE 3887 | p416-TEF-His-HGT1-HA with L370A mutation | Lab stock |
| **pP371A** | ABE 2853 | p416-TEF-His-HGT1-HA with P371A mutation | Lab stock |
| **pC372A** | ABE 3888 | p416-TEF-His-HGT1-HA with C372A mutation | Lab stock |
| **pL373A** | ABE 3888 | p416-TEF-His-HGT1-HA with L373A mutation | Lab stock |
| **pY374A** | ABE 3889 | p416-TEF-His-HGT1-HA with Y374A mutation | Lab stock |
| **TMD7** |  |  |  |
| **pS427A** | ABE 3662 | p416-TEF-His-HGT1-HA with S427A mutation | Lab stock |
| **pY428A** | ABE 3840 | p416-TEF-His-HGT1-HA with Y428A mutation | Lab stock |
| **pL429A** | ABE 3953 | p416-TEF-His-HGT1-HA with L429A mutation | Lab stock |
| **pL430A** | ABE 3924 | p416-TEF-His-HGT1-HA with L430A mutation | Lab stock |
| **pS431A** | ABE 3866 | p416-TEF-His-HGT1-HA with S431A mutation | Lab stock |
| **pY432A** | ABE 3841 | p416-TEF-His-HGT1-HA with Y432A mutation | Lab stock |
| **pA433G** | ABE 3842 | p416-TEF-His-HGT1-HA with A433G mutation | Lab stock |
| **pL434A** | ABE 3843 | p416-TEF-His-HGT1-HA with L434A mutation | Lab stock |
| **pN435A** | ABE 2005 | p416-TEF-His-HGT1-HA with N435A mutation (Kaur et al., 2009) | Lab stock |
| **pV436A** | ABE 3867 | p416-TEF-His-HGT1-HA with V436A mutation | Lab stock |
| **pA437G** | ABE 3954 | p416-TEF-His-HGT1-HA with A437G mutation | Lab stock |
| **pA438G** | ABE 3868 | p416-TEF-His-HGT1-HA with A438G mutation | Lab stock |
| **pV439A** | ABE 3844 | p416-TEF-His-HGT1-HA with V439A mutation | Lab stock |
| **pI440A** | ABE 3845 | p416-TEF-His-HGT1-HA with I440A mutation | Lab stock |
| **pA441G** | ABE 3846 | p416-TEF-His-HGT1-HA with A441G mutation | Lab stock |
| **pV442A** | ABE 4009 | p416-TEF-His-HGT1-HA with V442A mutation | Lab stock |
| **pV444A** | ABE 3515 | p416-TEF-His-HGT1-HA with V444A mutation | Lab stock |
| **pH445A** | ABE 2235 | p416-TEF-His-HGT1-HA with H445A mutation (Kaur et al., 2009) | Lab stock |
| **pC446A** | ABE 4060 | p416-TEF-His-HGT1-HA with C446A mutation | Lab stock |
| **pI447A** | ABE 4012 | p416-TEF-His-HGT1-HA with I447A mutation | Lab stock |
| **pL448A** | ABE 3847 | p416-TEF-His-HGT1-HA with L448A mutation | Lab stock |
| **pY449A** | ABE 3848 | p416-TEF-His-HGT1-HA with Y449A mutation | Lab stock |
| **TMD9** |  |  |  |
| **A509G** | ABE 2643 | pTEF-His-HGT1-HA with A509G mutation in TMD9. | Lab stock |
| **W510A** | ABE 2644 | pTEF-His-HGT1-HA with W510A mutation in TMD9. | Lab stock |
| **A511G** | ABE 2645 | pTEF-His-HGT1-HA with A511G mutation in TMD9. | Lab stock |
| **F512A** | ABE 2646 | pTEF-His-HGT1-HA with F512A mutation in TMD9. | Lab stock |
| **A513G** | ABE 2647 | pTEF-His-HGT1-HA with A513G mutation in TMD9. | Lab stock |
| **W514A** | ABE 2648 | pTEF-His-HGT1-HA with W514A mutation in TMD9. | Lab stock |
| **A515G** | ABE 2649 | pTEF-His-HGT1-HA with A515G mutation in TMD9. | Lab stock |
| **F516A** | ABE 2650 | pTEF-His-HGT1-HA with F516A mutation in TMD9. | Lab stock |
| **L517A** | ABE 2651 | pTEF-His-HGT1-HA with L517A mutation in TMD9. | Lab stock |
| **I518A** | ABE 2652 | pTEF-His-HGT1-HA with I518A mutation in TMD9. | Lab stock |
| **S519A** | ABE 2653 | pTEF-His-HGT1-HA with S519A mutation in TMD9. | Lab stock |
| **L520A** | ABE 2654 | pTEF-His-HGT1-HA with L520A mutation in TMD9. | Lab stock |
| **V521A** | ABE 2655 | pTEF-His-HGT1-HA with V521A mutation in TMD9. | Lab stock |
| **N522A** | ABE 2656 | pTEF-His-HGT1-HA with N522A mutation in TMD9. | Lab stock |
| **F523A** | ABE 2657 | pTEF-His-HGT1-HA with F523A mutation in TMD9. | Lab stock |
| **I524A** | ABE 2658 | pTEF-His-HGT1-HA with I524A mutation in TMD9. | Lab stock |
| **P525A** | ABE 2659 | pTEF-His-HGT1-HA with P525A mutation in TMD9. | Lab stock |
| **Q526E** | ABE 2660 | pTEF-His-HGT1-HA with Q526E mutation in TMD9. | Lab stock |
| **I528A** | ABE 2661 | pTEF-His-HGT1-HA with I528A mutation in TMD9. | Lab stock |
| **L529A** | ABE 2662 | pTEF-His-HGT1-HA with L529A mutation in TMD9. | Lab stock |
| **TMD13** |  |  |  |
| **pM739A** | ABE 3957 | p416-TEF-His-HGT1-HA with M739A mutation | Lab stock |
| **pG740A** | ABE 3869 | p416-TEF-His-HGT1-HA with G740A mutation | Lab stock |
| **pA741G** | ABE 3958 | p416-TEF-His-HGT1-HA with A741G mutation | Lab stock |
| **pV743A** | ABE 3871 | p416-TEF-His-HGT1-HA with V743A mutation | Lab stock |
| **pA745G** | ABE 3872 | p416-TEF-His-HGT1-HA with A745G mutation | Lab stock |
| **pG746A** | ABE 3873 | p416-TEF-His-HGT1-HA with G746A mutation | Lab stock |
| **pV747A** | ABE 3874 | p416-TEF-His-HGT1-HA with V747A mutation | Lab stock |
| **pA748G** | ABE 3874 | p416-TEF-His-HGT1-HA with A748G mutation | Lab stock |
| **pI749A** | ABE 3959 | p416-TEF-His-HGT1-HA with I749A mutation | Lab stock |
| **pS750A** | ABE 3960 | p416-TEF-His-HGT1-HA with S750A mutation | Lab stock |
| **pV751A** | ABE 3961 | p416-TEF-His-HGT1-HA with V751A mutation | Lab stock |
| **pI753A** | ABE 3877 | p416-TEF-His-HGT1-HA with I753A mutation | Lab stock |
| **pI745A** | ABE 3961 | p416-TEF-His-HGT1-HA with pI745A mutation | Lab stock |
| **pF755A** | ABE 3878 | p416-TEF-His-HGT1-HA with F755A mutation | Lab stock |
| **pL756A** | ABE 3879 | p416-TEF-His-HGT1-HA with L756A mutation | Lab stock |
| **pC757A** | ABE 4062 | p416-TEF-His-HGT1-HA with C757A mutation | Lab stock |
| **pV758A** | ABE 3880 | p416-TEF-His-HGT1-HA with V758A mutation | Lab stock |
| **pQ759A** | ABE 3083 | p416-TEF-His-HGT1-HA with Q759A mutation | Lab stock |

**Supplementary Table S7.** List of oligonucleotides and their sequences used

| **Primer Name** | **Sequence (5' to 3')** |
| --- | --- |
| **Primers used for cloning** | |
| **RP_416CEN** | CTTCTGTTCGGAGATTACCGAATC |
| **FP_416CEN** | TCTAGAAAACTTAGATTAGATTGC |
| **FP_416Ura** | TCGCGCGTTTCGGTGATGAC |
| **RP_416Ura** | CTCGAGTCATGTAATTAGTTATGT |
| **FP_AtOPT1_Frag** | AGAAAGAAAGCATAGCAATCTAATCTAAGTTTTCTAGAGGATCCATGACGAGCGTTTTCG |
| **RP_AtOPT1_Frag** | GTGAATGTAAGCGTGACATAACTAATTACATGACTCGAGTTAAAACACGGGACAACCTTC |
| **FP_AtOPT2_Frag** | TTAGAAAGAAAGCATAGCAATCTAATCTAAGTTTTCTAGAGGATCCATGGCTGCGATTGA |
| **RP_AtOPT2_Frag** | CGTGAATGTAAGCGTGACATAACTAATTACATGACTCGAGTTAAAAAACAGGACAACCAT |
| **FP_AtOPT3_Frag** | TTAGAAAGAAAGCATAGCAATCTAATCTAAGTTTTCTAGAGAATTCATGGACGCGGAGAA |
| **RP_AtOPT3_Frag** | CGTGAATGTAAGCGTGACATAACTAATTACATGActcgagTTAGAAAACGGGACAGCCTT |
| **FP_AtOPT4_Frag** | TAGAAAGAAAGCATAGCAATCTAATCTAAGTTTTCTAGAATGGCCACCGCCGACGAATTC |
| **RP_AtOPT4_Frag** | GTGAATGTAAGCGTGACATAACTAATTACATGACTCGAGTTATTTAACCGGACAACCATC |
| **FP_AtOPT5_Frag** | TTAGAAAGAAAGCATAGCAATCTAATCTAAGTTTTCTAGAGGATCCATGGTAGGCTCTCT |
| **RP_AtOPT5_Frag** | CGTGAATGTAAGCGTGACATAACTAATTACATGACTCGAGTTAGAACACCGGGCAGCCCT |
| **FP_AtOPT6_Frag** | TTAGAAAGAAAGCATAGCAATCTAATCTAAGTTTTCTAGAGGATCCATGGGAGAGATAGC |
| **RP_AtOPT6_Frag** | CGTGAATGTAAGCGTGACATAACTAATTACATGACTCGAGCTAGAAGACGGGACAGCCTT |
| **FP_AtOPT7_Frag** | TTAGAAAGAAAGCATAGCAATCTAATCTAAGTTTTCTAGAGGATCCATGGAAGAATCAGA |
| **RP_AtOPT7_Frag** | CGTGAATGTAAGCGTGACATAACTAATTACATGACTCGAGTTACGTATAAAGCGGGCATC |
| **FP_AtOPT8_Frag** | TTAGAAAGAAAGCATAGCAATCTAATCTAAGTTTTCTAGAGGATCCATGAAAGACTTTAC |
| **RP_AtOPT8_Frag** | GTGAATGTAAGCGTGACATAACTAATTACATGACTCGAGTTAGAAAACAGGACAACCATG |
| **FP_Fol2_OH_BamHI** | ATATATGGATCCCGCATGAAAATTTAGGATTTGG |
| **RP_Fol2_OH_XhoI** | TATATACTCGAGGCCTTTCTCTATGGAGATTTCTC |
| **FP_fol2cas_EcoRI** | ATATATGAATTCTGGCCTCCTCTAGTACACTC |
| **RP_Fol2cas_EcoRI** | ATATATGAATTCGCAGCTTTAAATAATCGG |
| **Primers used for deletion** | |
| **FP_opt_delcas_Leu** | TCGTTAAGAATATTGACGAGGACGTCAATAATCTCACTGCGAACTGTGGGAATACTCAGG |
| **RP_opt_delcas_Leu** | TACCACCTGGGTACTGTACACACAAGAAGATGATGACGACCCTCCTTTTTCTCCTTCTTG |
| **Primers used for confirming deletion** | |
| **FP_opt_delcas_ck** | ATGAGTACCATTTATAGGGAG |
| **RP_opt1_delcas_ck** | TTACCACCATTTATCATAACC |
| **Primers used for sequencing and verification** | |
| **TEF_F** | TTGATATTTAAGTTAATAAACGG |
| **TEF_R** | TTCAGGTTGTCTAACTCCTTC |
